# Manganese mediates antiviral effects by driving an ATM-TBK1 phosphorylation signaling pathway

**DOI:** 10.1101/2025.08.20.671272

**Authors:** Hongyan Sui, Rosana Wiscovitch-Russo, Silvia Cachaco, Jun Yang, Whitney Bruchey, Sylvain laverdure, Qian Chen, Tomozumi Imamichi

## Abstract

We previously reported that manganese (Mn) enhances innate immune responses to viral infection by inducing phosphorylation of TANK-binding kinase 1 (TBK1) in an Ataxia-telangiectasia mutated (ATM)-dependent manner. However, the underlying mechanism by which how Mn induces TBK1 phosphorylation remained unclear. Here, we show that Mn dose-dependently induced TBK1 phosphorylation in the presence of ATM across multiple cell lines, as well as in primary human macrophages and T cells. This phosphorylation was abolished in ATM-deficient cells, and we identified cytoplasmic ATM as a key mediator. Immunoprecipitation assays revealed that Mn promoted ATM phosphorylation at Ser1893, Ser1981, and Ser2996. TBK1 interacted with phosphorylated ATM at early stages, but upon phosphorylation, TBK1 dissociated from the ATM–TBK1 complex. This dissociation coincided with enhanced antiviral cytokine production. Furthermore, Mn dose-dependently suppressed HIV replication by inducing multiple antiviral host factors and cytokines. Together, these findings identify a cytoplasmic ATM–TBK1 phosphorylation cycle as a critical regulator of antiviral innate immunity and suggest Mn supplementation as a potential therapeutic approach against HIV and other viral infections.

Graphical Abstract:
Mn-dependent activation of the ATM-TBK1 phosphorylation signaling pathway. At early time points, Mn phosphorylated ATM at multiple site. And then TBK1 bound to p-ATM and became phosphorylated. Phosphorylated TBK1 then dissociated from the complex at later stages (right panel). P-TBK1 participated in activating downstream TBK1-IRF signaling, thereby enhancing antiviral cytokines induction to DNA or RNA virus infection (left panel).

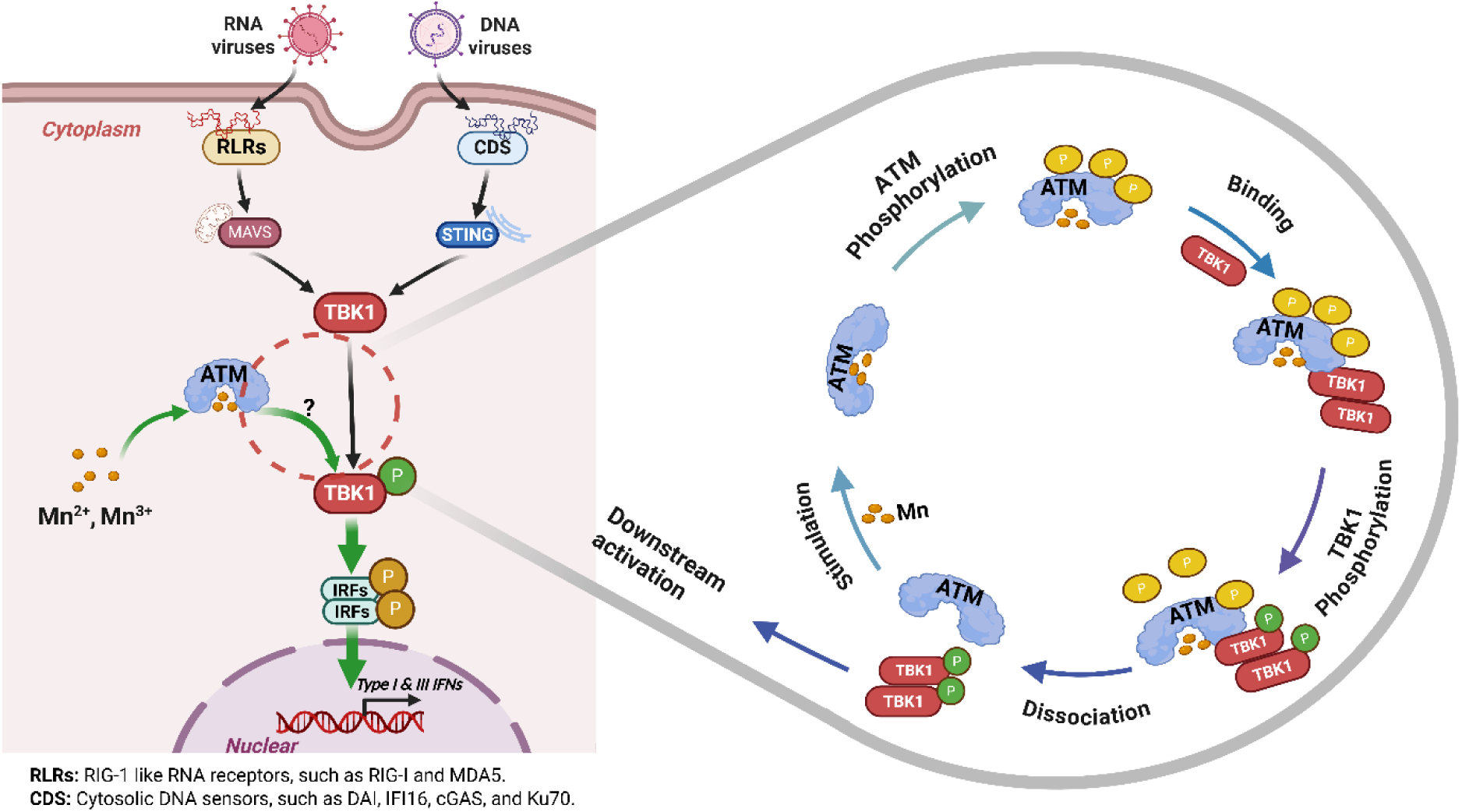

## Introduction

The innate immune system serves as the first line of host defense, aiming to prevent infection and eliminate invading pathogens such as viruses and bacteria (Medzhitov & Janeway, 2000; Pradeu *et al*, 2024). A key component of antiviral immunity involves pattern recognition receptors (PRRs), which detect conserved molecular features of viral pathogens and initiate signaling pathways that lead to the expression of antiviral genes (Akira *et al*, 2006). PRRs that detect extracellular pathogen-associated molecular patterns (PAMPs) are typically located on the plasma membrane or within endosomal membranes (Chan & Gack, 2016; Roudaire *et al*, 2020). These include Toll-like receptors (TLRs) (Akira *et al*, 2001; Delneste *et al*, 2007; West *et al*, 2006) and C-type lectin receptors (CLRs) (Geijtenbeek & Gringhuis, 2009; Reis e Sousa *et al*, 2024). Membrane-bound PRRs are primarily expressed in immune cells such as macrophages and dendritic cells (Sharma *et al*, 2025). In contrast, intracellular PRRs are found in the cytoplasm or nucleus of mammalian cells (Chan & Gack, 2016; Thompson *et al*, 2011). These include NOD-like receptors (NLRs) (Chen *et al*, 2009; Franchi *et al*, 2009), RIG-I-like receptors (RLRs) (Rehwinkel & Gack, 2020), and a group of intracellular DNA sensors such as cyclic GMP– AMP synthase (cGAS) (Cai *et al*, 2014; Sun *et al*, 2013; Yu & Liu, 2021) and interferon-γ (IFNγ)-inducible protein 16 (IFI16) (Unterholzner *et al*, 2010), some DNA repair proteins like Ku70 (Sui *et al*, 2021; Sui *et al*, 2017; Zhang *et al*, 2011) and DNA-PKcs (DNA-dependent protein kinase catalytic subunit) (Ferguson *et al*, 2012; Hristova *et al*, 2024). They play a crucial role in both DNA repair and innate immunity, particularly in the context of DNA sensing and signaling. These intracellular sensors are broadly or ubiquitously expressed across various cell types, enabling the detection of viral pathogens that have invaded the cytoplasm or nucleus of host cells.

Following the recognition of PAMPs, PRRs initiate innate immune signaling through the hierarchical activation of PRR family-specific adaptor proteins—for instance, mitochondrial antiviral-signaling protein (MAVS) (Bender *et al*, 2015; Kumar *et al*, 2006; Seth *et al*, 2005; Xu *et al*, 2005), stimulator of interferon genes (STING) (Ishikawa & Barber, 2011; Sui *et al*., 2017), and myeloid differentiation primary response 88 (MYD88) (Deguine & Barton, 2014; Medzhitov *et al*, 1998)—as well as a shared set of well-characterized serine/threonine kinases, which include TANK-binding kinase 1 (TBK1), IκB kinase (IKK) complex and IKK-related kinases: IKKε, and transcription factors: IRF1, IRF3, IRF7, nuclear factor kappa B (NF-kB) (Iwanaszko & Kimmel, 2015; Pham & TenOever, 2010; Sato *et al*, 2000; tenOever Benjamin *et al*, 2004). This signal transduction cascade ultimately results in the production of various host defense molecules, including type I and type III interferons (IFNs), along with pro-inflammatory cytokines and chemokines. The secreted IFNs act in both autocrine and paracrine manners by binding to their respective receptors, thereby inducing the expression of hundreds of interferon-stimulated genes (ISGs). The proteins encoded by ISGs inhibit critical steps in the viral life cycle and modulate innate immune sensing and cytokine production, contributing to the establishment of an antiviral state (Dalskov *et al*, 2023; Katze *et al*, 2002).

Despite rapid advances in the development of antiviral drugs for some viruses, there remains a concerning lack of effective antiviral drugs for many clinically significant viral pathogens. The continued emergence of new and previously known viral pathogens, as well as drug-resistant variants, highlights the continuous and urgent need for the development of novel and more effective vaccines and antivirals to combat viruses and mitigate human disease. For example, HIV has evolved multiple sophisticated strategies to evade recognition by innate immune sensors and to suppress the activation of PRRs and their downstream signaling pathways, thereby posing significant challenges to the development of effective preventive and therapeutic interventions (Fang *et al*, 2025; Mouzakis *et al*, 2025).

Mn is an important dietary trace element associated with several physiological processes such as anti-tumor immunity, development and bone growth (Chen *et al*, 2018; Li & Yang, 2018). In addition to its well-established role in general health, Mn has recently attracted attention for its involvement in regulating innate immune responses. Mn was shown to increase the sensitivity of cGAS to double-stranded DNA (dsDNA) and its enzymatic activity also facilitates STING activity by boosting cGAMP-STING binding affinity (Wang *et al*, 2018). Further studies have revealed that Mn directly activates cGAS to induce a noncanonical catalytic synthesis of 2′3′-cGAMP, through a conformation closely resembling that of dsDNA-activated cGAS (Sun *et al*, 2023). Mn has also been shown to activate natural killer (NK) cells in neonatal mice through promoting the level of IFN (Ming *et al*, 2024). In addition, the human blood-derived monocytes/macrophages exposed to Mn show an increased production of the cytokines: interleukin (IL)-1β, IL-6, IL-8, IFN-γ and tumor necrosis factor α (TNF-α) as compared to control-treated cells (Filipov *et al*, 2005). Consistent with those studies, we found that Mn enhances the induction of multiple antiviral cytokines—including IFN-α, IFN-β, and IFN-λ—but by promoting the phosphorylation of TBK1 (Sui *et al*, 2022). TBK1 is a serine/threonine protein kinase that plays a central role in multiple signaling pathways, acting as a critical regulatory node (Louis *et al*, 2018). Due to its involvement in diverse cellular processes, selectively modulating TBK1 activity within specific pathways offers a more effective strategy for targeted regulation. In addition, ATM is also a serine/threonine protein kinase, and is primarily activated in response to DNA double-strand breaks and orchestrates a complex DNA damage response by phosphorylating a wide range of substrates, including key tumor suppressors such as p53 (Biddlestone-Thorpe *et al*, 2013; Karakostis *et al*, 2024). This activation leads to cell cycle arrest, DNA repair, or apoptosis, depending on the extent of the damage (Stracker *et al*, 2013). We previously demonstrated that ATM is involved in the Mn-enhanced innate immune signaling pathway (Sui *et al*., 2022). However, the precise mechanism by which Mn promotes TBK1 phosphorylation through ATM remains unclear, and the role of ATM in innate immunity is still not fully understood. Therefore, the present study aims to provide new insights into the regulatory function of Mn in antiviral immunity, particularly in the context of HIV and other viral infections.

## Results

### Mn dose-dependently induces phosphorylation of TBK1 across multiple cell types with the presence of functional ATM

In our previous study, we demonstrated that Mn enhances DNA-mediated IFN induction in human primary macrophages and showed that ATM plays a role in this Mn-enhanced innate immune response, as evidenced by ATM knockdown and treatment with ATM inhibitors (Sui *et al*., 2022). To address potential off-target effects of siRNA and the non-specificity of ATM inhibitors, we employed ATM KO A549 cells to further validate the essential role of ATM. Compared to wild-type A549 cells (A549 WT), phosphorylation of TBK1 was markedly reduced in ATM KO A549 cells (Figure 1A). Furthermore, Mn-induced enhancement of IFN-λ1 expression was completely abolished in ATM KO A549 cells (Figure 1B). These results indicated that Mn induced TBK1 phosphorylation in an ATM-dependent manner. To determine whether these observations extend beyond A549 cells, we next evaluated other cell lines and human primary cells. 293T cells, HeLa cells, human primary monocyte-derived macrophages (MDMs), and PHA-activated human primary CD4(+) T cells were included in the study. The results demonstrated that Mn induced phosphorylation of TBK1 in a dose-dependent manner across all these cell types, with ATM expression confirmed in each cell type (Figures 1C, 1D, 1E, and 1F). Notably, the optimal Mn concentration varied among cell types: for example, phosphorylation of TBK1 in macrophages was effectively induced at 25 µM Mn (Figure 1E), whereas HeLa cells required a higher concentration of 500 µM for optimal induction (Figure 1D). Overall, data from Figure 1 indicated that primary cells respond to lower Mn concentrations compared to established cell lines. Collectively, these findings confirmed that ATM was essential for Mn-induced phosphorylation of TBK1 across diverse cell types.

**Figure 1.**
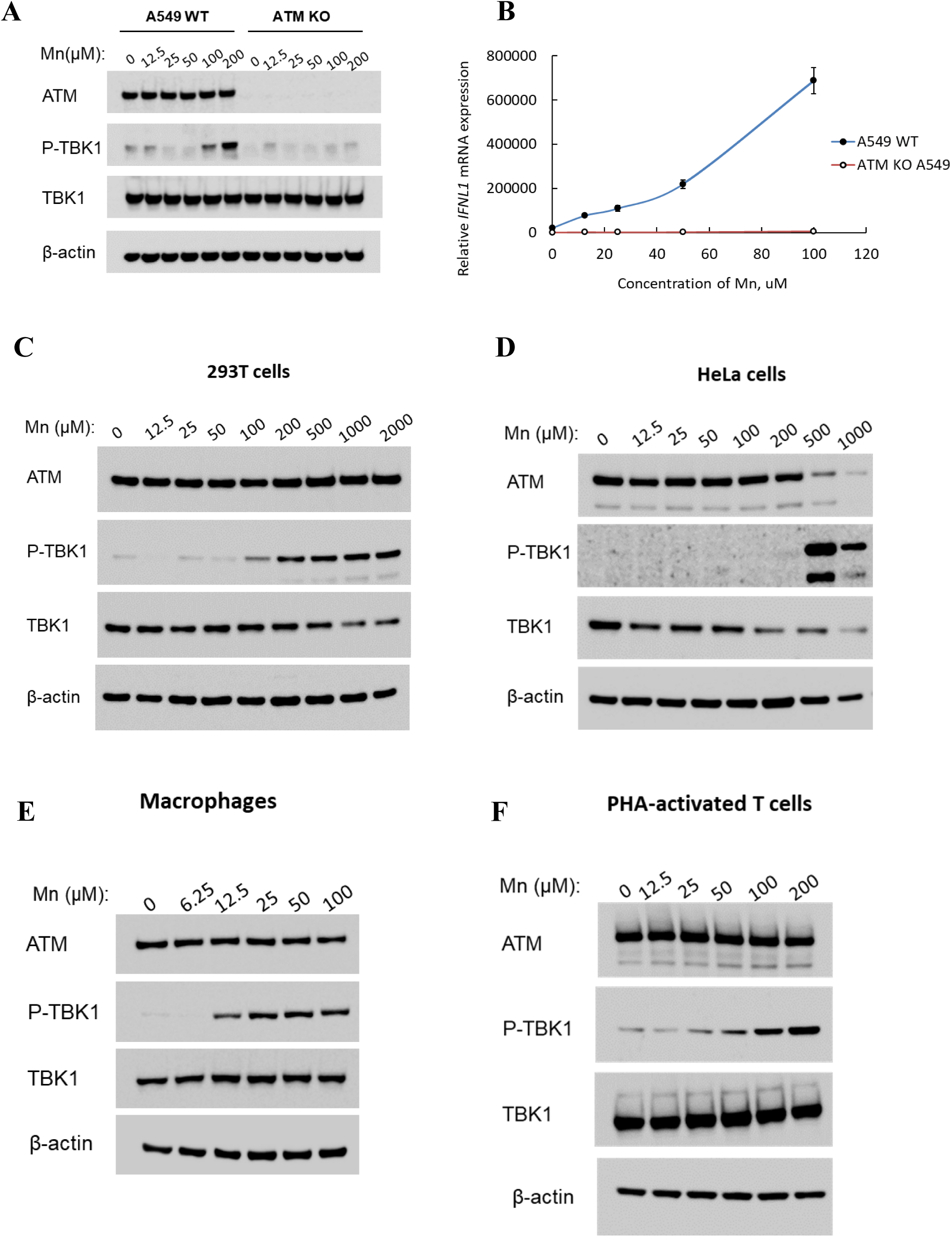
Mn dose-dependently phosphorylates TBK1 in multiple cell types in an ATM-dependent manner. **(A)** A549 wild-type and ATM KO A549 cells were treated with increasing concentrations of MnCL_2_. Total cell lysates were collected 24 hours post-treatment using RIPA buffer and analyzed by Western blot using antibodies against ATM, TBK1, and p-TBK1. β-actin served as a loading control. **(B)** A549 and ATM KO A549 cells were treated with the indicated concentrations of MnCL_2_, followed by stimulation with linearized DNA 24 hours later. After an additional 24 hours, total RNA was extracted, and IFN-λ1 mRNA expression levels were measured by quantitative RT-PCR and normalized to GAPDH. **(C–F)** 293T cells (**C**), HeLa cells (**D**), MDMs (**E**), and PHA-activated CD4⁺ T cells (**F**) were treated with varying doses of MnCL_2_. Whole cell lysates were collected after 24 hours and analyzed by Western blot using anti-ATM, anti-p-TBK1, and anti-TBK1 antibodies. β-actin was included as an internal loading control.

### Cytoplasmic ATM promotes phosphorylation of TBK1

To define whether the interaction between ATM and TBK1 occurred, we performed immunofluorescence analysis using confocal microscopy. Mn-treated A549 cells were collected at 6, 14, and 24 h post-treatment. Cells were fixed and stained with anti-ATM and anti-phospho-TBK1 antibodies, then visualized under a confocal microscope. Confocal microscopy images revealed that ATM localized both in the nucleus and the cytoplasm of A549 cells (Figure 2A). However, its nuclear expression was notably more intense, indicating that ATM is predominantly a nuclear kinase. In contrast, the phosphorylation of TBK1 progressively increased with the duration of Mn treatment. By 24 hours post-treatment, a substantial increase in phosphorylated TBK1 levels was observed, characterized by a distinct punctate pattern within the cytoplasm. This suggested that Mn induced cytoplasmic activation of TBK1 over time. The merged images further suggested that cytoplasm-localized ATM may interact with TBK1 and potentially contribute to the induction of TBK1 phosphorylation.

**Figure 2.**
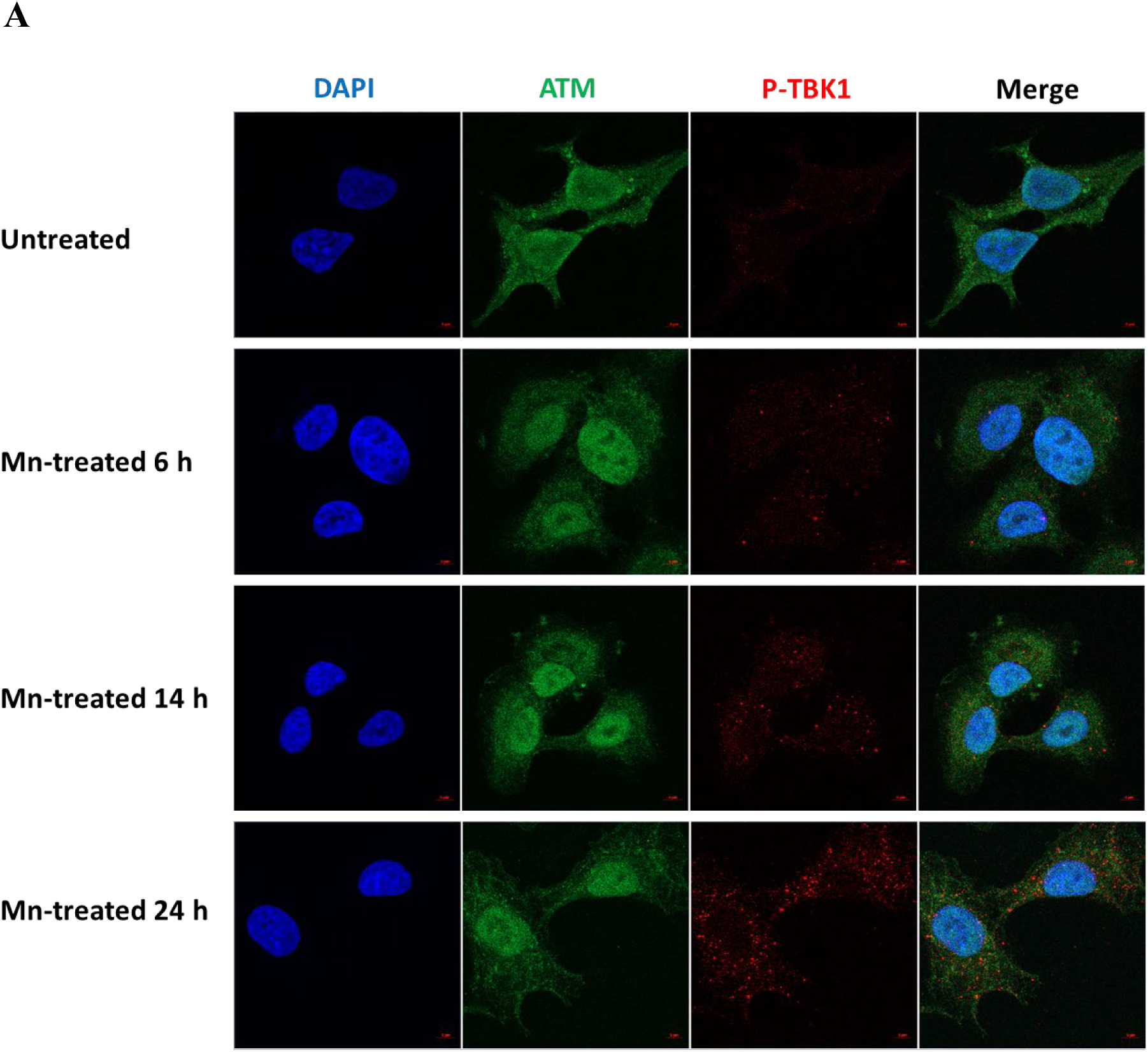
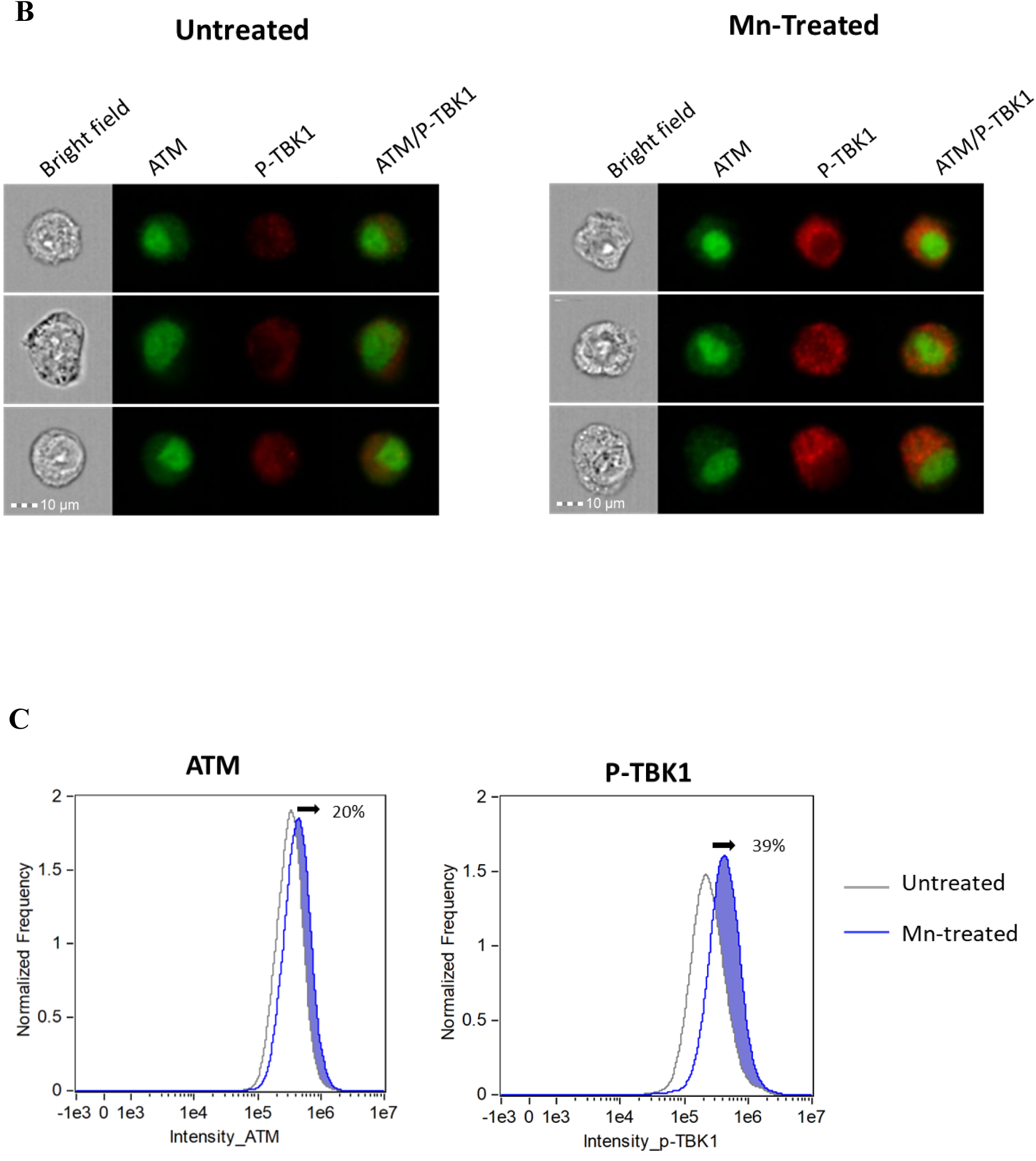
Cytoplasmic ATM promotes phosphorylation of TBK1. **(A)** Confocal microscopy analysis of the localization and expression of ATM and p-TBK1 in A549 cells after Mn treatment. A549 cells were grown on 12mm coverslip-inserted 12-well cell culture plates and treated with or without 100 µM MnCL_2_, then fixed, permeabilized and stained with anti-ATM (green), anti-P-TBK1 (Red) antibody at 6, 14 and 24 h after Mn treatment. Nuclei were counterstained with DAPI (blue), and cells were imaged using a 63×/1.4 objective. **(B-C)** AMINS analysis for the expression of ATM and p-TBK1. A549 cells were seeded on 6-well cell culture plates and treated with or without Mn treatment at 100 µM, then detached using Accutase at 24 h after Mn treatment, fixed, permeabilized, and stained by anti-ATM (Green) and anti-p-TBK1 (Red) antibodies, the single cell morphonology was viewed by bright field. **(B)** Image galleries display of brightfield, fluorescence dyes AF647 (ATM, channel 11) and AF488 (p-TBK1, channel 2), and composite image of the two fluorescence channels. **(C)** Histogram overlay of ATM and p-TBK1 intensity of untreated versus Mn-treated focused single cells. The shifted area was shaded in blue, and the shifted percentage (calculated by Fiji) were indicated in the figure.

To further quantify Mn-induced phosphorylation of TBK1 (p-TBk1) and assess the subcellular localization of phosphorylated TBK1 in relation to ATM, Amnis ImageStream flow cytometry was performed. Approximately 50,000 focused cells were acquired for each sample. Single-color controls and isotype antibody controls were included to generate a spectral crosstalk matrix, which was applied to each dataset for accurate spectral compensation across detection channels. Representative single-cell images were shown in Figure 2B. These images demonstrated that ATM expression levels and localization remained largely unchanged between Mn-treated and untreated cells. In contrast, p-TBK1 expression was markedly increased in Mn-treated cells compared to untreated controls, indicating robust induction of TBK1 phosphorylation following Mn exposure. To further quantify the ImageStream data, the intensity distributions—defined as the sum of all pixel intensities per cell—for ATM and p-TBK1 were analyzed and displayed as histograms for both untreated and Mn-treated subpopulations. Histogram overlays were used to compare the fluorescence intensity profiles between the two groups. As shown in Figure 2C, the Mn-treated cell population exhibited an approximate 20% increase in overall ATM intensity compared to the untreated group. Notably, around 40% of the Mn-treated cells shifted outside the distribution range of the untreated population, indicating a substantial increase in p-TBK1 expression following 24 h Mn exposure. The isotype control staining for ATM or p-TBK1 was shown in the supplementary Figure 1. These findings, consistent with the confocal immunofluorescence results, further confirmed that Mn significantly enhanced the expression of p-TBK1 in the cytoplasm. Based on these data, we hypothesized that the cytoplasm serves as the primary site of interaction between cytoplasmic TBK1 and cytoplasm-localized ATM.

### TBK1 initially interacts with ATM and then dissociates from the ATM–TBK1 complex upon its own phosphorylation

To further investigate the mechanism of the interaction between ATM and TBK1 in the cytoplasm, a co-immunoprecipitation (co-IP) assay was performed to determine whether ATM and TBK1 formed a protein complex. First, ATM KO 293T cells were transfected with a FLAG-tagged ATM plasmid and treated with or without Mn. Cell lysates were collected at 6 and 14 hours of post-treatment, followed by immunoprecipitation using anti-Flag antibodies to pull down ATM and any interacting proteins. Endogenous TBK1 was detected in the ATM-containing protein complex 6 hours after Mn treatment. Notably, TBK1 formed a complex under untreated conditions, with similar band intensities observed regardless of Mn treatment. This suggested that TBK1 constitutively interacted with ATM, independent of Mn exposure, at early time points. However, analysis of immunoprecipitated samples collected at 14 hours post-Mn treatment showed a slight reduction in TBK1 band intensity in the Mn-treated samples compared to untreated controls (Figure 3A). Based on these results, we hypothesized that TBK1 bound to ATM at early time points, and following its phosphorylation, phosphorylated TBK1 dissociated from the ATM–TBK1 complex at later stages. To further validate this hypothesis, an additional immunoprecipitation assay was performed using a model overexpressing both ATM and TBK1. Cell lysates were collected 14 hours after Mn treatment. Consistent with previous observations, the intensity of the TBK1 band was clearly reduced in Mn-treated samples compared to untreated controls (Figure 3B). In summary, the immunoprecipitation assays demonstrated that TBK1 associated with ATM at early time points and subsequently dissociated from the complex following its own phosphorylation at later stages.

**Figure 3.**
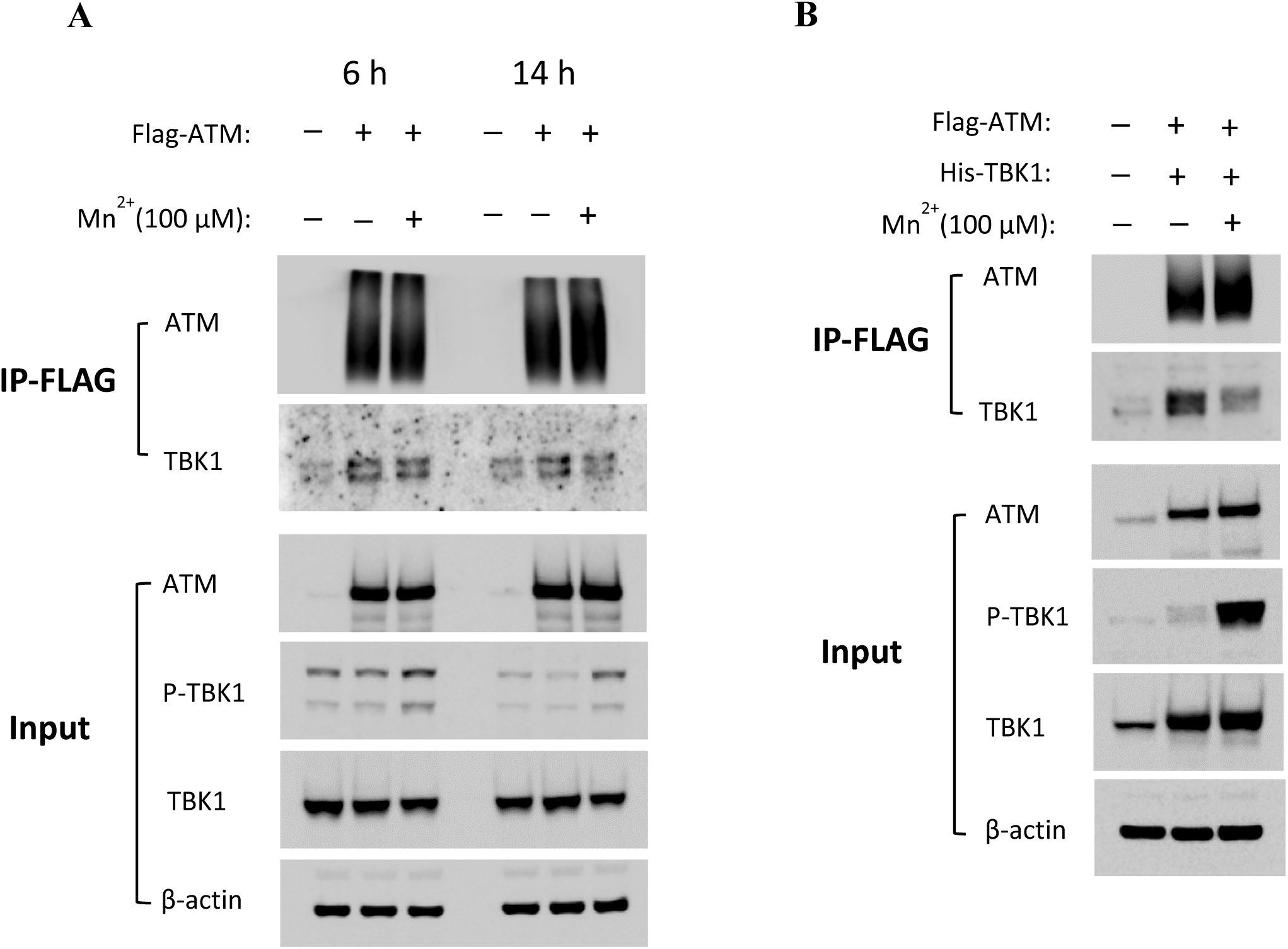
TBK1 first interacts with ATM and later dissociates from the ATM–TBK1 complex after it becomes phosphorylated. ATM KO 293T cells were transfected with **(A)** Flag-tagged ATM or **(B)** a combination of plasmids encoding Flag-tagged ATM and His-tagged TBK1 and then were treated by MnCL_2_ at 100 µM. Cytosolic lysates were collected at 6 or 14 h after Mn treatment, followed by immunoprecipitation (IP) with anti-Flag antibody-conjugated agarose. Precipitated proteins were then analyzed by Western blotting with antibodies against ATM and TBK1. Input controls were also included.

### Mn induces phosphorylation of ATM at multiple sites at early time points, followed by subsequent phosphorylation of TBK1 at later stages

While we have demonstrated that Mn treatment induced phosphorylation of TBK1, it remained unclear whether ATM itself undergoes phosphorylation in response to Mn, and what the precise relationship is between phosphorylated ATM and TBK1 phosphorylation. PTM-MS analysis was performed on FLAG-immunoprecipitated samples from Mn-treated ATM KO 293T cells overexpressing FLAG-tagged ATM. The phosphorylation status of ATM was examined, revealing that Mn treatment induced phosphorylation at multiple sites—Ser1893, Ser1981, and Ser2996—each located within distinct ATM kinase domains (Figure 4A). These findings confirmed that Mn triggered multiple-site phosphorylation of ATM (Figure 4A). To validate the PTM-MS findings, Western blot analysis was performed using an anti-phospho-ATM (Ser1981) antibody to assess ATM phosphorylation. Phosphorylation of TBK1 was also examined. The results, shown in Figure 4B, demonstrated that Mn-treated samples exhibited a stronger p-ATM (Ser1981) signal compared to untreated controls. Quantification of band intensity using Image J analysis (right panel in Figure 4B) further confirmed the significant increase in ATM phosphorylation upon Mn treatment. In comparison, the band intensity of p-TBK1 in Mn-treated cell lysates was significantly stronger than in untreated samples. These data suggested that Mn first induced ATM phosphorylation, which in turn promoted the phosphorylation of TBK1.

**Figure 4.**
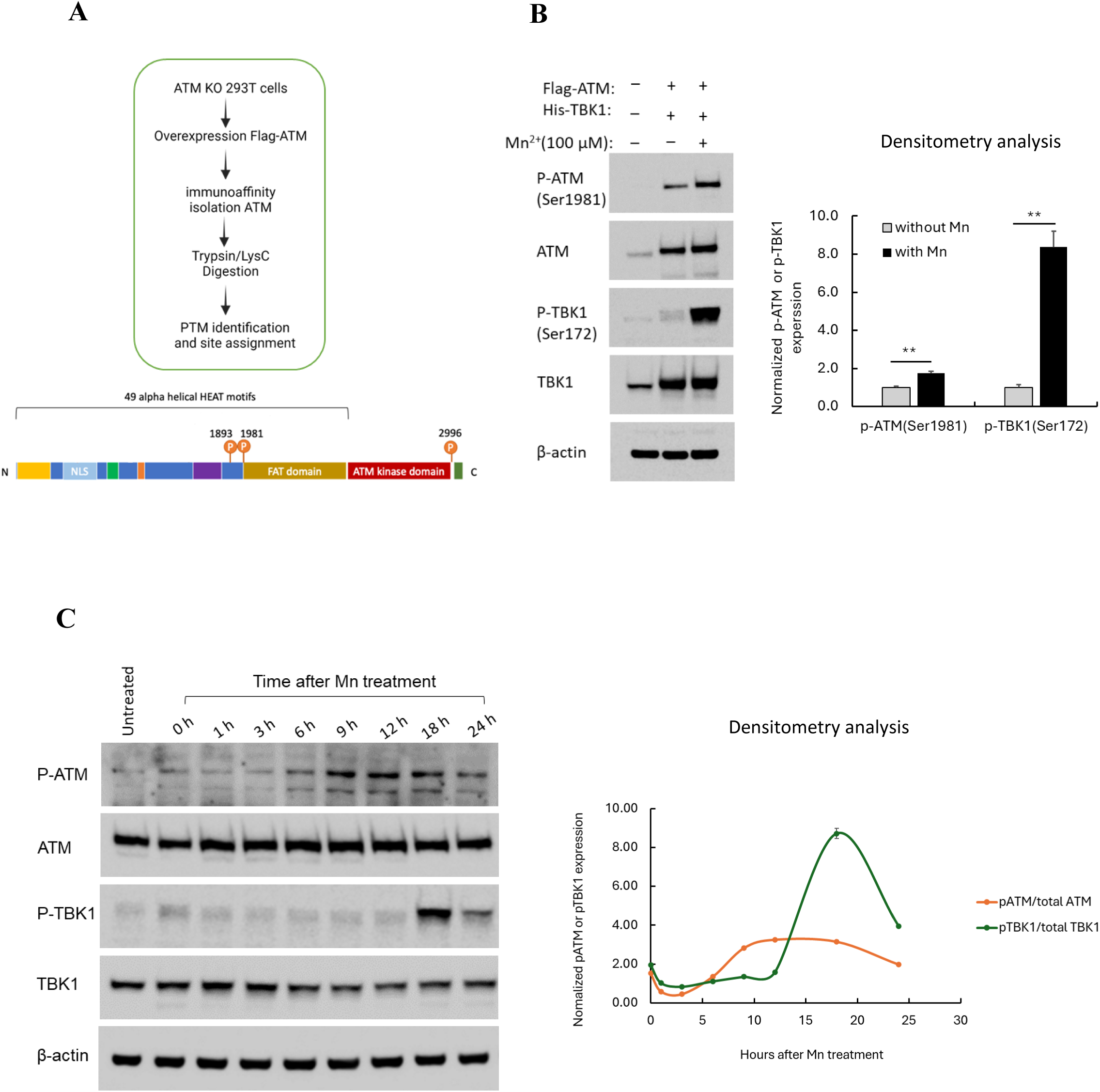
Mn phosphorylates ATM at multiple sites at an earlier time point. **(A)** The illustration for Mass spectrometry process and identified phosphorylated sites on ATM by Mn treatment. **(B)** ATM KO 293T cells were overexpressed by Flag-tagged ATM and His-tagged TBK1 and then treated by Mn at 100 µM. The whole cell lysate was prepared and analyzed by western blot using the antibodies against p-ATM (Ser1981), ATM, p-TBK1 (Ser172), anti-β-actin was included to indicate the loading control (left panel). A densitometry analysis was performed according to the data shown in the left panel by Fiji (right panel). **(C)** A549 cells were seeded in 6-well cell culture plates and treated with Mn at 100 µM. The total cell lysate was prepared using RIPA buffer at different time points after Mn treatment. The cell lysate was analyzed by anti-p-ATM, anti-ATM, anti-p-TBK1 and TBK1 antibodies. Anti-β-actin was included as the internal loading control for the experiment. A densitometry analysis was performed according to the data shown in left panel by Fiji, the intensity of band for p-ATM or p-TBK1 is normalized by total ATM or TBK1, respectively (right panel).

To confirm the temporal sequence of Mn-induced phosphorylation of ATM and TBK1, Western blot analysis was performed on samples collected at various time points following Mn treatment. Phosphorylation of ATM was detected using an anti-phospho-ATM (Ser1981) antibody, while phosphorylation of TBK1 was assessed with an anti-phospho-TBK1 (Ser172) antibody. The data showed that ATM phosphorylation began at 6 hours post-Mn treatment, peaked at 12 hours, and then gradually declined. In contrast, phosphorylation of TBK1 was first detected at 18 hours after Mn treatment, followed by a subsequent decrease in signal intensity (Figure 4C, left panel). The data clearly demonstrated that ATM phosphorylation occurred earlier, between approximately 9 to 18 hours after Mn treatment, while TBK1 phosphorylation started later, around 18 hours post-treatment. The dynamic phosphorylation cascade from ATM to TBK1 was further demonstrated by the densitometric analysis presented in the right panel of Figure 4C. This temporal pattern suggested that Mn first induced ATM phosphorylation, which was then followed by phosphorylation of TBK1. Accordingly, we proposed that Mn triggered a phosphorylation cascade involving ATM activation upstream of TBK1.

### One single mutation within kinase domain or any other phosphorylation sites of ATM has no substantial impact on Mn-induced TBK1 phosphorylation

Mn induced phosphorylation of ATM at multiple sites—Ser1893, Ser1981, and Ser2996—each located within distinct domains of the ATM kinase protein (Figure 4A). To determine which domain or phosphorylation site of ATM is critical for the subsequent phosphorylation of TBK1, various ATM mutants were transfected into ATM KO 293T cells. These ATM mutant-expressing cells were then compared to wild-type 293T cells as well as ATM KO 293T cells reconstituted with wild-type ATM. Consistently, in 293T cells expressing endogenous ATM, Mn treatment induced TBK1 phosphorylation in a dose-dependent manner. In contrast, TBK1 phosphorylation was markedly reduced in ATM KO 293T cells. Notably, a detectable level of phosphorylated TBK1 was only observed at a high Mn concentration (200 µM) in the absence of ATM as Mn-dependent autophosphorylation. Reintroduction of wild-type ATM into ATM KO cells fully restored Mn-induced TBK1 phosphorylation (the third panel, Figure 5). However, overexpression of kinase-dead ATM or ATM carrying a mutation at Ser1981 still resulted in partial recovery of TBK1 phosphorylation (the fourth and fifth panel, Figure 5), indicating that ATM kinase activity and phosphorylation at multiple sites were critical for full activation of TBK1 by Mn. These data suggested that phosphorylation at multiple sites on ATM collectively contributed to Mn-induced TBK1 phosphorylation. Loss of phosphorylation at any single site did not significantly impair TBK1 activation, indicating a redundant or cooperative mechanism among these phosphorylation sites.

**Figure 5.**
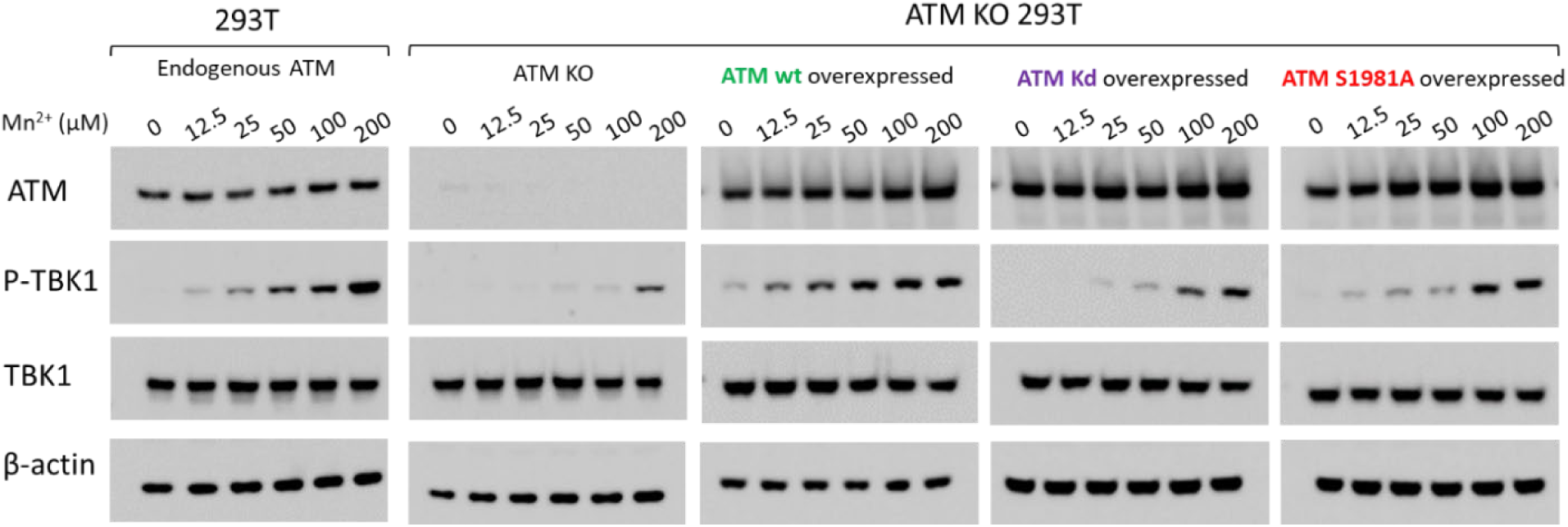
One single mutation within kinase domain or any other phosphorylation sites of ATM has no substantial impact on Mn-induced TBK1 phosphorylation. ATM KO 293T cells were overexpressed by transfecting plasmids encoding wild type ATM (wt ATM), mutated ATM at kinase domain (ATM kd), mutated ATM at S1981A (ATM S1981A), respectively, and treated with Mn at varying concentrations. The total cell lysate was collected at 24 h after Mn treatment and subjected to western blot analysis using anti-ATM, p-TBK1, TBK1 antibodies, anti-β-actin was included as internal loading control. Untransfected 293T cells were also included in the experiment to serve as a positive control.

### Mn inhibits HIV replication through the induction of multiple antiviral cytokines and factors with anti-HIV activity

Our previous study has demonstrated that Mn inhibits both DNA and RNA virus infections (Sui *et al*., 2022). The current data further indicated the broad-spectrum antiviral potential of Mn, suggesting its capacity to suppress a wide range of viral infections. Given that HIV remains a major global public health concern, we aimed to investigate whether Mn treatment can also inhibit HIV replication in human primary cells. Human primary macrophages were pretreated with varying concentrations of Mn prior to infection with pseudotyped HIV Luc-V. The results demonstrated that Mn inhibited HIV replication in a dose-dependent manner (Figure 6A).

**Figure 6.**
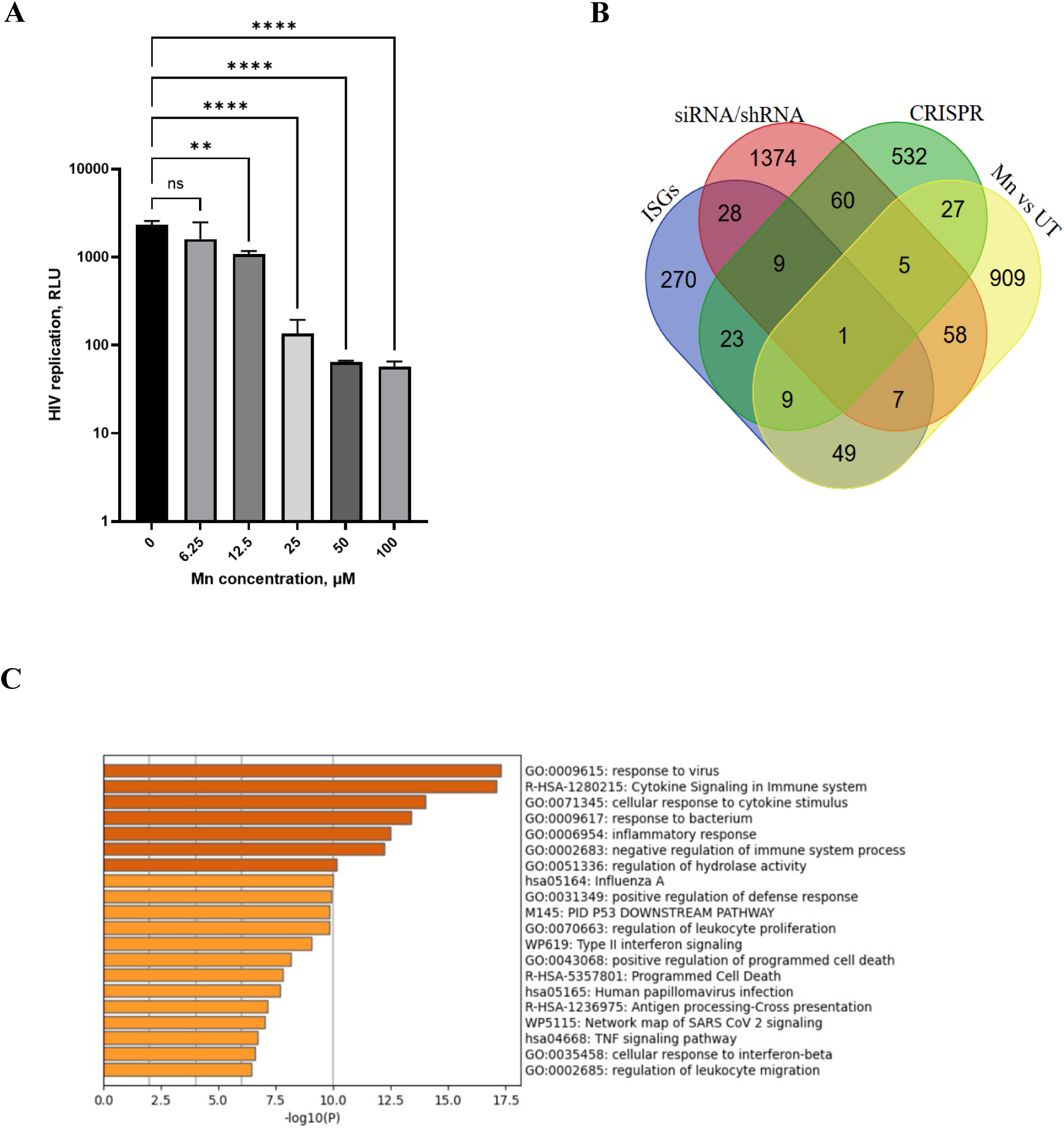
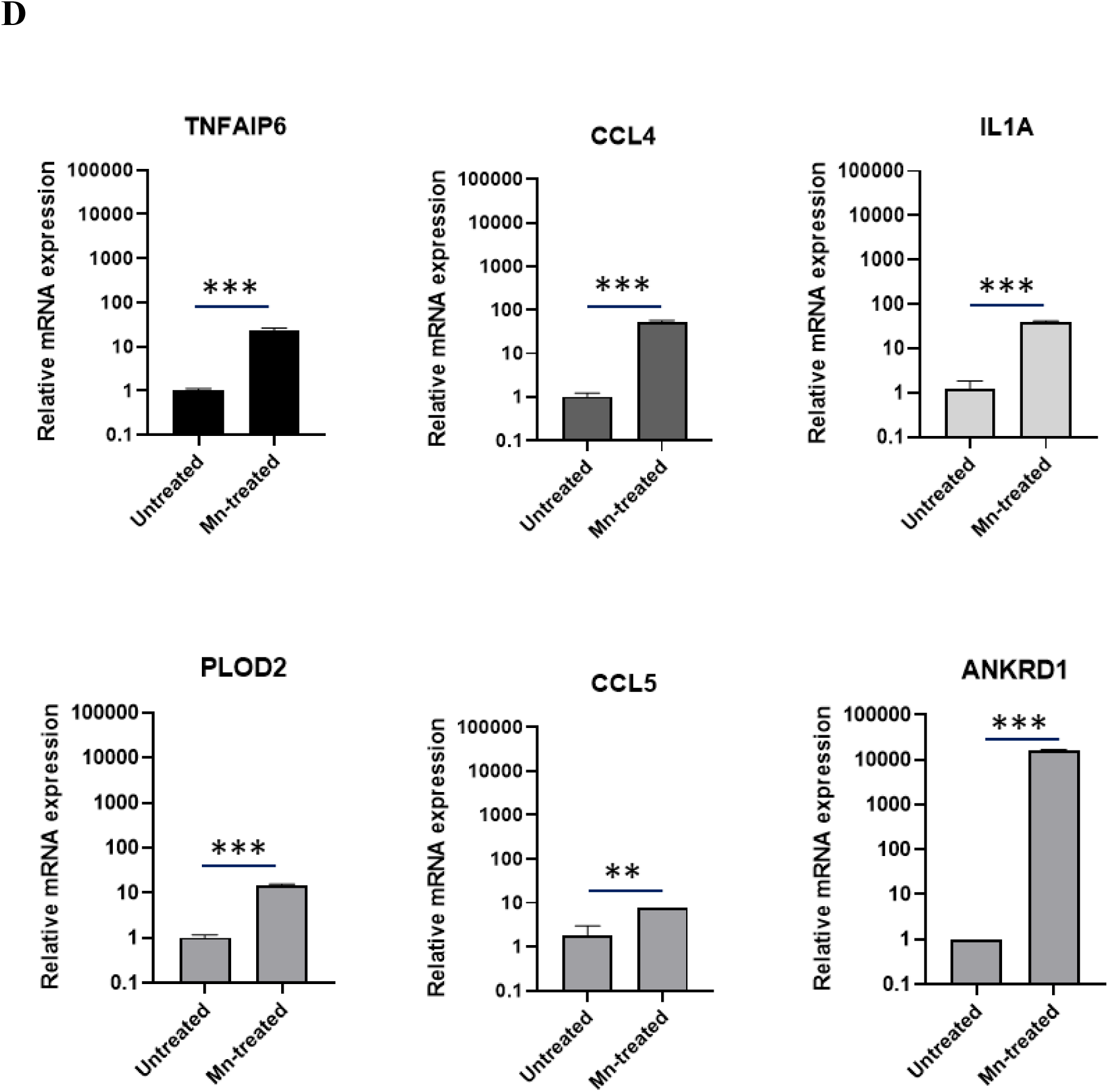
Mn inhibits HIV virus replication in MDMs by inducing multiple anti-viral cytokines or ISGs. **(A)** MDMs were treated with different concentration of MnCL_2_, and 24 h later, were further infected with recombinant HIV-Luc virus. HIV replication was determined luciferase assay at 48 h after infection. One-way ANOVA was performed to indicate significant changes among samples by P value in different dose of Mn treatment vs. untreated control. **<0.01, ****<0.0001, and ns (not significant) were noted. **(B)** Identification of host factors associated with HIV inhibition in the Mn-induced 1065 genes. Venn diagram analysis was conducted using the Mn-induced 1065 genes and a total of 2,439 genes of host factor identified from CRISPR, siRNA/shRNA library screening, and ISGs. **(C)** Functional enrichment analysis by Metascape (Metascape.org) for 156 Mn-induced genes related with anti-HIV activities (Supplementary Table S1). **(D)** Total RNA was extracted from with or without Mn-treated MDMs. Relative mRNA expression of indicated genes was detected using real-time RT-PCR, the values were normalized by GAPDH expression. Student’s *t* test was performed to indicate significant changes among samples by P-value between Mn-treated and untreated samples, respectively. **<0.01 and ***<0.001 were noted.

Notably, Mn treatment at a concentration of 25 µM resulted in over 90% inhibition of viral replication, suggesting that Mn may induce some factors with potent anti-HIV activities. To identify host genes induced by Mn treatment, we performed microarray analysis comparing gene expression profiles in human primary macrophages with and without Mn pretreatment. This analysis revealed 1,065 genes that were upregulated or downregulated by more than 2-fold with a p-value < 0.05. To assess the potential relevance of these genes to HIV inhibition, we conducted a Venn diagram analysis comparing the 1,065 of Mn-induced genes with a dataset of 2,439 known anti-HIV host factors identified through CRISPR, siRNA, and shRNA library screenings, as well as interferon-stimulated genes (ISGs) (Kane *et al*, 2016; König *et al*, 2008; Krasnopolsky *et al*, 2020; Li *et al*, 2020; OhAinle *et al*, 2018; Park *et al*, 2017; Zhou *et al*, 2008) (Figure 6B). Interestingly, 42 of the Mn-induced genes overlapped with anti-HIV genes identified via CRISPR screening, 71 were identified through siRNA/shRNA library screenings, and 66 were present in the ISG database. In total, 156 of the 1,065 of Mn-induced genes overlapped with previously identified anti-HIV host factors, suggesting that Mn treatment may activate a network of antiviral genes that contribute to the suppression of HIV replication. To further characterize the 156 of Mn-induced potential anti-HIV genes in MDMs, functional annotation analysis was performed using Metascape (Zhou *et al*, 2019). This analysis revealed significant enrichment of pathways related to viral response, cytokine signaling in the immune system, and cellular response to cytokine stimulus (Figure 6C). And the list of 156 of Mn upregulated or downregulated anti-HIV factors was shown in the Supplementary Table 2. To validate the expression of selected antiviral genes, real-time RT-PCR was conducted on Mn-treated MDMs. Specifically, the expression levels of TNFAIP6, CCL4, IL1A, PLOD2, CCL5, and ANKRD1 were measured using gene-specific probes. The results confirmed that all six genes were significantly upregulated following Mn treatment (Figure 6D), which is consistent with the microarray data.

## Discussion

We have previously reported that supplementation of Mn in the form of Mn^2+^ in culture medium enhances innate antiviral immune response against DNA or RNA virus infection. This phenomenon is due to an increase in the phosphorylation of TBK-1. The use of siRNA against ATM or Ku-60019 (Golding *et al*, 2009; Shu *et al*, 2023), an ATM inhibitor demonstrated that ATM, a protein kinase, is involved in this pathway (Sui *et al*., 2022). However, both siRNA and small molecule inhibitors can produce unintended side effects. siRNA, even when designed for specific mRNA silencing, may inadvertently suppress transcripts with similar sequences or trigger unexpected signaling pathways (Ali Zaidi *et al*, 2023). Conversely, small-molecule inhibitors can engage off-target proteins or pathways, leading to unforeseen biological effects (Weiss *et al*, 2007). Therefore, whether ATM is directly involved in the Mn-mediated enhancement of the innate immune response remains to be fully confirmed, and the molecular mechanisms underlying the interaction network among Mn, ATM, and TBK1 have not yet to be elucidated. In this study, we aimed to fully elucidate the role of ATM in the Mn-induced TBK1 phosphorylation pathway. Experiments using ATM KO A549 and ATM KO 293T cells consistently demonstrated that both TBK1 phosphorylation and the Mn-mediated enhancement of the antiviral innate immune response were abolished in the absence of ATM. These findings provide further evidence that ATM is involved in Mn-induced TBK1 phosphorylation.

ATM is a versatile protein kinase with diverse roles in DNA damage response, cellular homeostasis, and disease pathogenesis (Bhatti *et al*, 2011; Ueno *et al*, 2022). It is considered as a nuclear protein associated with the chromatin and nuclear matrix (Gately *et al*, 1998). We demonstrated that ATM localizes to both the nucleus and cytoplasm, indicating that it may also exert functional roles in the cytoplasm. We further confirmed that p-TBK1 is predominantly localized in the cytoplasm, and its expression intensity progressively increased with the duration of Mn exposure. Based on this observation, we subsequently demonstrated that cytoplasmic ATM interacts with cytoplasmic TBK1. Moreover, the punctate staining pattern of p-TBK1 suggests that TBK1 forms homodimers upon phosphorylation (Larabi *et al*, 2013).

In the immunoprecipitation assay, the amount of TBK1 bound to ATM decreased at 14 h post-treatment compared with samples collected at the same time points but without Mn treatment. This observation suggested that unphosphorylated TBK1 interacted with ATM at the initial stage, and that the TBK1–ATM interaction is a transient process. TBK1 dissociated from the ATM–TBK1 complex after phosphorylation, illustrating the dynamic nature of the ATM– TBK1 interaction. Subsequently, phosphorylated TBK1 may translocate to other regions of the cytoplasm to activate downstream signaling pathways. This process is consistent with the biology function of TBK1 as a kinase protein in innate immune response (Suzuki *et al*, 2013).

TBK1 is a kinase involved in innate immunity, autophagy, and cell cycle regulation. It can be found in various cellular locations, including the cytosol, endosome membrane, and nucleoplasm. Interestingly, TBK1’s localization is often regulated by interactions with adaptor proteins, which can direct it to specific cellular compartments and signaling pathways (Paul *et al*, 2025; Runde *et al*, 2022; Wang *et al*, 2024). So, the translocation of TBK1 within the cytoplasm is consistent with the function switches of TBK1.

ATM is a large protein kinase, a member of the phosphatidylinositol 3-kinase-like protein kinase (PIKK) family (Bhatti *et al*., 2011). ATM is known as a master regulator of the cellular response to DNA double-strand breaks (DSBs) and related genotoxic stress in the nucleus (Shibata & Jeggo, 2021). When DSBs occur, ATM becomes an activated form and then initiates a complex signaling network by phosphorylating various downstream proteins, including p53, Chk2, and H2AX, leading to processes like cell cycle arrest and DNA repair (Cao *et al*, 2006; Karakostis *et al*., 2024). In the current study, we demonstrated ATM also phosphorylated TBK1, and ATM facilitated TBK1-mediated activating of downstream signaling. However, what is the status of ATM after Mn treatment? Mn is an essential trace element and serves as a cofactor for many enzymes (Li & Yang, 2018). Studies show that Mn can activate ATM kinase activity (Paull, 2015; Tidball et al., 2015). A specific signaling pathway involving ATM and the tumor suppressor protein p53 has been identified to be Mn dependent (Tidball *et al*, 2015). Our current study suggested that Mn phosphorylated ATM at multiple sites, Ser1893, Ser1981, and Ser2996, and all the phosphorylation sites contributes to downstream TBK1 phosphorylation, since single mutation at one phosphorylation site had no critical effect on the activity of TBK1 phosphorylation. However, in the context of DNA damage, autophosphorylation at Ser1981 is widely recognized as a hallmark of ATM activation. Phosphorylation at this site stabilizes ATM at DSBs and is essential for an effective DNA damage response (So *et al*, 2009). And further we confirmed that the phosphorylation of ATM is the upstream event of TBK1 phosphorylation.

Inactive ATM typically forms a homodimer. This dimeric structure is crucial for maintaining ATM in an autoinhibited state. In this form, key regulatory elements within the ATM dimer restrict access to the active site, preventing substrate binding and kinase activity (Lau *et al*, 2016; Ueno *et al*., 2022; Warren & Pavletich, 2022). Upon activation, the ATM dimer dissociates, forming active monomers. The monomeric form allows for a structural rearrangement that removes the inhibitory constraints on the active site. This liberation of the active site allows ATM to bind and phosphorylate its target proteins (Du *et al*, 2014). Active ATM then phosphorylates a variety of downstream proteins involved in DNA repair, cell cycle control, and apoptosis. For example, ATM phosphorylates members of the MRN (MRE11-RAD50-NBS1) complex, which are crucial for repairing DNA damage (Phan & Rezaeian, 2021). And we confirmed in the current study that Mn initially phosphorylated ATM, and the active form of ATM then phosphorylated TBK1, followed by an enhanced induction of innate immune response. Thus, the Mn-mediated ATM-TBK1 phosphorylation is essential for Mn-mediated antiviral activities.

In this study, we delineated an Mn-enhanced antiviral innate immune response mediated by an ATM–TBK1 phosphorylation axis, wherein ATM activation positively correlated with TBK1 phosphorylation. ATM exhibits an Mn-dependent function, a previously unrecognized role in innate immune responses that are commonly present in multiple cell types. The ATM–TBK1 dynamic cycle is essential for the Mn-mediated innate immune response. However, Zhang et al. reported an alternative ATM–TBK1 model, where ATM inhibition triggers TBK1 activation and type I IFN production in pancreatic cells, a response that is further amplified by radiation (Zhang *et al*, 2019). In their study, radiation-induced DNA damage leads to the release of damaged DNA into the cytoplasm, which activates the cGAS–STING pathway, culminating in TBK1 activation and IFN production. In this pathway, ATM does not directly phosphorylate TBK1; rather, ATM inhibition reduces phosphorylation of CHK2 and AKT2 while increasing phosphorylation of p70-S6K and SRC, with phosphorylated SRC subsequently phosphorylating TBK1 (Hu & Zhang, 2024). Therefore, we speculated that the interplay between ATM and TBK1 is stimulus– and cell-type dependent, and the signaling pathway may differ under distinct biological contexts.

Wang *et al*. reported that Mn binds to cGAS, increasing its sensitivity to dsDNA and its production of the secondary messenger cGAMP. Mn increases the binding affinity between cGAMP and STING. Activating the cGAS-STING pathway leads to the production of type I interferons (IFNs) and other cytokines (Wang *et al*., 2018). Mn was also shown essential in the innate immune sensing of tumors (Lv *et al*, 2020; Ma *et al*, 2025). Mn administration promotes the maturation of dendritic cells and macrophages and enhances tumor-specific antigen presentation (Wang *et al*., 2018; Zhang *et al*, 2021). Our previous study revealed that Mn enhances both DNA and RNA virus-mediated antiviral activities through increasing the phosphorylation of TBK1 (Sui *et al*., 2022), implicating that Mn may work in any TBK1-involved signaling pathway. We further evaluated the potential antiviral activity of Mn on HIV virus infection and the data suggested that Mn dose-dependently inhibits HIV virus infection in human primary macrophages. This data further motivated us to investigate whether Mn induces some antiviral factors which has been validated with antiviral activities previously. The microarray analysis indicated that 156 of the 1,065 Mn-induced genes overlapped with previously identified anti-HIV host factors (Huang *et al*, 2019; Kane *et al*., 2016; König *et al*., 2008; Krasnopolsky *et al*., 2020; Li *et al*., 2020; OhAinle *et al*., 2018; Park *et al*., 2017; Taylor *et al*, 2018; Zhou *et al*., 2008). And those genes were highly focus on the response to virus or bacterium, cytokine signaling in immune system. So, we proposed that Mn treatment may activate a network of antiviral factors that contribute to the suppression of HIV replication. In the current study, we used recombinant HIV virus express the Vesicular Stomatitis Virus glycoprotein (VSV-G) instead of HIV envelop protein, so the viruses enter macrophages without using HIV receptor (CD4 and chemokine receptors, CCR5 or CXCR4) and complete only a single round of infection. As such, this system allowed us to focus on HIV-1 infection at a post-entry level (Borok *et al*, 2001; Cronin *et al*, 2005). We have planned to use replication competent HIV virus for infection in next step to further fully evaluate the antiviral efficiency of Mn with more physiology relevance. However, the current finding provided valuable insight on developing new anti-HIV reagents to fight HIV virus infection on humans. Since Mn induced 156 of multiple anti-HIV factors to suppress HIV virus infection, it is supposed there would be less opportunities for HIV virus develop Mn-resistant mutations. The study focusing on how Mn inhibits HIV infection is ongoing, the finding from which will provide more details regarding the mechanism and feasibility.

In summary, we further elucidated the role of ATM in Mn-mediated antiviral responses. Mn first induced phosphorylation of ATM, which in turn phosphorylated TBK1. Initially, TBK1 interacted with monomeric ATM and became phosphorylated. Upon phosphorylation, TBK1 dissociated from the ATM complex and activated downstream signaling pathways, thereby enhancing the antiviral response. The broad-spectrum antiviral activity of Mn also inhibits HIV replication, highlighting its great potential for the development of novel anti-HIV therapeutic formulations.

## Material and methods

Approval for these studies, including all sample materials, was granted by the National Institute of Allergy and Infectious Diseases Institutional Review Board, and participants gave written informed consent prior to blood being drawn. All experimental procedures in these studies were approved by the National Cancer Institute at Frederick (the protocol code number: 2016– 19A6/11-30) and performed in accordance with the relevant guidelines and regulations.

### Cells

SV40 T-antigen-transformed Human embryonic kidney 293 (293T) and HeLa cells were obtained from the American Type Culture Collection (ATCC, Manassas, VA, USA) and maintained according to the manufacturer’s instructions in D10 medium (DMEM medium (Thermo Fisher Scientific, Waltham, MA, USA) supplemented with 10% fetal bovine serum (FBS; R&D systems, Minneapolis, MN, USA), 25 mM HEPES (Quality Biology, Gaithersburg, MD, USA), and 5 µg/mL gentamicin (Thermo Fisher Scientific)). A549 cells (ATCC) and ATM knockout (KO) A549 cells (Ubigene, Austin, TX, USA) were cultured in F-12K medium (Thermo Fisher Scientific, Waltham, MA, USA) supplemented with 10% FBS (R&D systems), 25 mM HEPES (Quality Biology), and 5 µg/mL gentamicin (Thermo Fisher Scientific). ATM KO 293T cells (Ubigene) were cultured in D10 medium as described above.

To generate monocyte-derived macrophages (MDMs), CD14^+^ monocytes were isolated from peripheral blood mononuclear cells (PBMCs) of healthy donors using CD14^+^ microbeads (Miltenyi Biotec, San Diego, CA) and plated at 1.0 × 10^7^ cells per 10 cm petri dish (Thermo Fisher Scientific), and then differentiated into MDMs by stimulation with 25 ng/mL M-CSF (R&D Systems) in macrophage serum-free medium (Thermo Fisher Scientific) for seven days. Differentiated MDMs were then maintained in D10 medium prior to experimental use.

CD4⁺ T cells were isolated from CD14^-^ PBMCs of healthy donors (the flow-through from prior CD14⁺ cell isolation) using CD4⁺ microbeads (Miltenyi Biotec), following the manufacturer’s protocol (Miltenyi Biotec). Isolated cells were stimulated with 5 μg/mL phytohemagglutinin (PHA; Sigma-Aldrich, St. Louis, MO, USA) for three days in RPMI-1640 medium (Thermo Fisher Scientific) supplemented with 10% FBS (Thermo Fisher Scientific), 25 mM HEPES (Quality Biological), and 5 μg/mL gentamicin (Thermo Fisher Scientific).

### Plasmid and DNA transfection

The plasmids pcDNA3.1(+) Flag-His-ATM wild-type (wt), pcDNA3.1(+) Flag-His-ATM kinase-dead (kd), and hATM-S1981A (Addgene, Watertown, MA, USA) were used to overexpress either wild-type or mutant forms of ATM. The plasmid PEF1α-V5-His-TBK1, constructed from the empty vector pEF1α (Thermo Fisher Scientific), was used to overexpress TBK1 for co-immunoprecipitation assays. Transfections into 293T or ATM KO 293T cells were performed using TransIT-293 transfection reagent (Mirus Bio, Madison, WI, USA) according to the manufacturer’s instructions. To generate linearized noncoding DNA for stimulation assays, the pCR2.1 plasmid (Thermo Fisher Scientific) was digested with EcoRI, followed by purification using a PCR purification kit (QIAGEN, Germantown, MD, USA). The resulting linearized DNA was used as a noncoding DNA stimulant, as previously described (Sui *et al*, 2013). DNA stimulation of MDMs, CD4^+^ T cells and A549 cells was carried out using TransIT-X2 transfection reagent (Mirus Bio) according to the manufacturer’s protocol.

### RNA extraction and real-time RT-PCR

Total cellular RNA was extracted using the RNeasy Mini Kit (QIAGEN) according to the manufacturer’s instructions. Complementary DNA (cDNA) was synthesized from total RNA using TaqMan Reverse Transcription Reagents (Thermo Fisher Scientific) with random hexamer primers (Thermo Fisher Scientific). Gene expression levels of IFNs or Mn-stimulated genes were quantified using the QuantStudio™ 7 Pro Real-Time PCR System (Thermo Fisher Scientific). The thermal cycling conditions consisted of 40 cycles of 95 °C for 15 seconds and 60 °C for one minute. Relative mRNA levels were calculated using the ΔΔCt method (Livak & Schmittgen, 2001), with GAPDH serving as the internal reference gene. Normalized expression was presented as the fold change relative to the average ΔCt value of the control group. TaqMan gene expression probes specific for IFNs or Mn-responsive genes were obtained from Applied Biosystems (Thermo Fisher Scientific).

### Western blot

Whole-cell lysates were prepared using RIPA buffer (Boston BioProducts, Ashland, MA, USA) supplemented with a protease inhibitor cocktail (Sigma-Aldrich) and Halt phosphatase inhibitor cocktail (Thermo Fisher Scientific). Protein concentrations were determined using the microBCA Protein Assay Kit (Thermo Fisher Scientific) to ensure equal loading across all samples. Equal amounts of total protein were resolved on either NuPAGE 4–12% Bis-Tris gels or 3–8% Tris-Acetate gels (Thermo Fisher Scientific), followed by transfer onto 0.45 µm nitrocellulose or PVDF membranes (Thermo Fisher Scientific). Membranes were probed with the appropriate primary antibodies, followed by HRP-conjugated secondary antibodies. Protein bands were visualized using ECL Plus Western blotting detection reagents (GE Healthcare, Chicago, IL, USA) and imaged on a Chemiluminescent Western Blot Imager Azure 300 (Azure Biosystems, Dublin, CA, USA). Primary antibodies and secondary antibodies are listed in the Supplementary table 1. The dilutions in the experiments are provided.

### Co-Immunoprecipitation assay and mass spectrometry analysis

ATM KO 293T cells were seeded at a density of 3.5 × 10⁶ cells per 10-cm dish and incubated overnight. Cells were then transfected with 6 µg of the following plasmids: pcDNA3.1(+) Flag-His-ATM (wild-type) (Addgene) and/or PEF6/V5-His-TBK1. After 48 hours, cells were treated with 100 µM MnCl₂ (Sigma-Aldrich). Cells were lysed 24 h after Mn treatment in Pierce IP Lysis Buffer (Thermo Fisher Scientific), supplemented with protease and phosphatase inhibitor cocktail (Sigma-Aldrich and Thermo Fisher Scientific). Lysates were incubated overnight at 4 °C with anti-FLAG M2 affinity gel (MilliporeSigma). Beads were washed four times with tris-buffered saline (TBS, Quality Biology), and bound proteins were eluted by boiling for 5 minutes at 95 °C in NuPAGE™ LDS Sample Buffer (Thermo Fisher Scientific). The resulting samples were analyzed by immunoblotting for ATM and TBK1 expression.

For proteomic analysis, FLAG-immunoprecipitated proteins from Mn-treated and untreated ATM KO 293T cells were subjected to trypsin/Lys-C digestion, followed by liquid chromatography–tandem mass spectrometry (LC-MS/MS). Raw MS data were analyzed using Proteome Discoverer 2.5 (Thermo Fisher Scientific) against the human protein database. Proteins identified with matched peptide sequences and post-translational modifications were cataloged for each condition, and a comparative analysis between Mn-treated and untreated samples was performed by PooChon Scientific (Frederick, MD, USA).

### Immunofluorescence assay and confocal microscopy

A549 cells were seeded at a density of 1.0 × 10^5^ cells per well onto Corning BioCoat Poly-L-Lysine–treated 12 mm glass coverslips (MilliporeSigma) in 12-well plates and treated with or without 100 µM MnCl₂. At 6, 14, and 24 hours post-treatment, cells were fixed with 4% formaldehyde (Thermo Fisher Scientific) for 15 minutes at room temperature. After fixation, cells were washed three times with 1X phosphate-buffered saline (PBS, Quality Biology), permeabilized with 0.1% Triton X-100 (Sigma-Aldrich) in PBS for 10 minutes, and washed again three times with PBS. Cells were then blocked with Intercept Blocking Buffer (LI-COR Biotechnology, Lincoln, NE, USA) for at least 1 hour at room temperature. Coverslips were removed from the wells and placed face-down on one 80 μL-drop of primary antibody solution (prepared in blocking buffer) and incubated overnight at 4 °C in a humidified dark chamber. The following day, coverslips were washed three times with PBS and incubated with appropriate fluorophore-conjugated secondary antibodies for 1 hour at room temperature, followed by four times of PBS washes. The information about the antibodies used in this assay are listed in the Supplementary Table S1. Finally, coverslips were mounted on glass slides using ProLong Diamond Antifade Mountant with DAPI (Thermo Fisher Scientific), and images were acquired using a Zeiss Axio Observer.Z1 LSM800 confocal microscope equipped with a Plan-Apochromat 63×/1.40 oil immersion objective.

### Imaging flow cytometry and data analysis

A549 cells were seeded at a density of 1.0 × 10^6^ cells per well in 6-well plates and treated with 100 µM MnCl₂. At 24 hours post-treatment, cells were harvested using Accutase (Thermo Fisher Scientific) and fixed with 4% formaldehyde (Thermo Fisher Scientific) in 1X PBS for 15 minutes at room temperature, followed by permeabilization with 0.1% Triton X-100 (Sigma-Aldrich) in 1x PBS for 10 minutes. Cells were incubated with primary antibodies (see immunofluorescence assay section) overnight at 4 °C, then washed and incubated with Alexa Fluor–conjugated secondary antibodies (Cell Signaling Technology)) for 1 hour at room temperature. After each incubation step, cells were washed three times with 1X PBS. Finally, cells were resuspended in 1× PBS containing 2% BSA (Sigma-Aldrich) for downstream analysis.

Stained cells were analyzed using the ImageStream® X Mark II Imaging Flow Cytometer (Amnis Corporation, Cytek Biosciences, Seattle, WA, USA) at 40× magnification under low flow rate, high sensitivity, and high gain settings. The following laser lines and detection parameters were used: 488 nm laser (40.00 mW) for AF488 detection in channel 2 (606–680 nm), 647 nm laser (150 mW) for AF647 detection in channel 11 (642–746 nm), 785 nm laser (2.66 mW) for side scatter (SSC) in channel 6 (746–800 nm); brightfield images were captured in channels 1 and 9, while fluorescence images were acquired in Channel 2: Phospho-TBK1 (AF488), Channel 11: ATM (AF647), and Channel 6: Side scatter respectively. A total of 50,000 single-cell events were acquired per sample. Single-cell gating was performed using brightfield images (channel 1) based on cell area and aspect ratio. Cells were gated to include those with an area between 200 and 600 with an aspect ratio (width-to-height) greater than 0.6, ensuring selection of intact, single cells and exclusion of doublets or debris.

The intracellular expression levels of phosphorylated TBK1 (p-TBK1) and ATM in untreated and Mn-treated cells were analyzed using IDEAS™ software (version 6.4). A specific subpopulation of focused, single cells was defined using the following gating strategy: Focus quality was determined by the Gradient RMS of the brightfield image (channel 1), with a threshold of >50 to identify in-focus cells. Single cells were then gated based on area (200–600) and aspect ratio (>0.6), using brightfield parameters, as described above. Visual inspection of image galleries confirmed the accuracy of gate placement for each individual sample. The fluorescence intensity distributions—measured as the sum of all pixel intensities per cell—for p-TBK1 (AF488, channel 2) and ATM (AF647, channel 11) were plotted as histograms for both untreated and Mn-treated cell populations. Histogram overlays were used to directly compare intensity distributions between conditions. To further examine the distribution of p-TBK1, bivariate plots were generated showing brightfield area vs. p-TBK1 intensity (channel 2). Isotype controls (AF488 and AF647) for both untreated and Mn-treated samples were used to define background fluorescence resulting from nonspecific antibody binding. The antibodies used for staining were Mouse anti-ATM (1:100 dilution) and Rabbit anti–p-TBK1 (1:50 dilution).

### Pseudotyped HIV-1 luciferase viruses and infection

VSV-G-pseudo typed HIV-luciferase virus (HIV Luc-V) was prepared by co-transfecting HEK293T cells with pNL4-3ΔEnv-Luc (Connor *et al*, 1995; He *et al*, 1995) and pLTR-VSVG (Chang *et al*, 1999) using the TransIT-293 transfection reagent (Mirus Bio) as previously described (Dai *et al*, 2013). Virus-containing supernatants of HEK293T cell were harvested 48 h after transfection and filtered with 0.45-μm Steriflip Filter Units (MilliporeSigma). Virus particles were pelleted by ultracentrifugation at 100,000×g for 2 h, at 4 °C on a 20% sucrose in 150 mM NaCl-HEPES, pH7.4 buffer and resuspended in D10 medium. Virus concentration was quantified using Alliance HIV-1 P24 antigen ELISA kit (Revvity, Boston, MA, USA), and then aliquoted, stored at –80 °C until use.

HIV-1 Infection assay was performed using the pseudotyped HIV Luc-V as previously described (Imamichi *et al*, 2023). Monocytes-derived macrophages were seeded in 96 well plate (50 × 10^3^ cells/well), and then infected with 50 μL of 100 ng p24/ml of HIV Luc-V for 2 h at 37 °C. Infected cells were washed twice with warm D10 medium and then cultured for 24 h at 37 °C. Cells were lysed in 1X Passive Lysis Buffer (Promega, Madison, Wisconsin, USA) and luciferase activity was measured using the Luciferase Assay System (Promega) on an Enspire Multimode Plate Reader (Perkin Elmer, Boston, MA, USA).

### Microarray

Microarray analysis was conducted as previously described (Sui *et al*., 2022). Briefly, total cellular RNA was extracted using the RNeasy isolation kit (QIAGEN). RNA quantity and quality were assessed using Nanodrop 1000 spectrophotometer (Thermo Fisher Scientific) and an Agilent Bioanalyzer RNA Nano 6000 chip (Agilent Technologies, Santa Clara, CA, USA). Complementary RNA (cRNA) synthesis, labeling, and hybridization to the Human GeneArray (Thermo Fisher Scientific) were performed according to the manufacturer’s instructions.

Raw expression data were processed and analyzed using Partek Genomics Suite (Partek, Inc., St. Louis, MO, USA). Arrays were normalized via quantile normalization, and statistically significant gene expression differences were identified using two-way ANOVA. Functional enrichment analyses were carried out using Metascape, a comprehensive resource for gene annotation, visualization, and integrated discovery.

### Statistical analysis

Results represent data from at least three independent experiments. Values are expressed as mean ± standard deviation (SD) of individual samples. The student’s unpaired *t*-test or one way ANOVA was used and p values lower than 0.05 were considered significant. The p-values with less than 0.05 is noted as *, less than 0.01 as **, less than 0.001 as ***, and less than 0.0001 as ****.

## Acknowledgments

We thank HC. Lane and M. Baseler for supporting this project. We also thank Ming Hao and Weizhong Chang for discussing protein structure analysis. The Graphical Abstract was created using Biorender.com. This project has been funded in whole or in part with federal funds from the National Cancer Institute, National Institutes of Health, under contract number HHSN261200800001E. This research was supported, in part, by the National Institute of Allergy and Infectious Diseases, National Institutes of Health. The content of this publication does not necessarily reflect the views or policies of the Department of Health and Human Services, nor does the mention of trade names, commercial products, or organizations imply endorsement by the U.S. Government.

## Disclosure and competing interests statement

The authors declare that they have no conflict of interest.

## Author contributions

H.S. designed experiments, performed assays, analyzed data and wrote the manuscript; R.W., S.C., W.B. and Q.C. conducted experiments; J.Y. and S.L. analyzed microarray and AMINS data, respectively; and T.I. created the project, oversaw the data, supervised the project and revised the manuscript.

## Supplementary information

**Supplementary Figure 1.** Sequential gating strategy for analysis in IDEAS™ (v6.4) analysis software. **(A)** Histogram of channel 1 brightfield (BF) gradient RMS of all acquired cells for selection of focused cells. **(B)** Selection of single cells from focused cells population using a density scatter plot of brightfield area versus brightfield aspect ratio. **(C-D)** Histogram overlay of intensity of focused single cell subpopulations of AF647 isotype control versus untreated and Mn-treated cells. **(E-F)** Histogram overlay of intensity of focused single cell subpopulations of AF488 isotype control versus untreated and Mn-treated cells.

**Supplementary Table 1.** A list of antibodies used for western blot, immunofluorescence and Imaging flow cytometry assays. The information about the catalog numbers, suppliers and dilutions are provided.

**Supplementary Table 2.** Gene list for Mn stimulated anti-HIV factors in MDMs (folds changes>2). This table is referred to Figure 6B&C.

## References

1. Akira S, Takeda K, Kaisho T (2001) Toll-like receptors: critical proteins linking innate and acquired immunity. Nat Immunol 2: 675–680

2. Akira S, Uematsu S, Takeuchi O (2006) Pathogen Recognition and Innate Immunity. Cell 124: 783–801

3. Ali Zaidi SS, Fatima F, Ali Zaidi SA, Zhou D, Deng W, Liu S (2023) Engineering siRNA therapeutics: challenges and strategies. Journal of Nanobiotechnology 21: 381

4. Bender S, Reuter A, Eberle F, Einhorn E, Binder M, Bartenschlager R (2015) Activation of Type I and III Interferon Response by Mitochondrial and Peroxisomal MAVS and Inhibition by Hepatitis C Virus. PLOS Pathogens 11: e1005264

5. Bhatti S, Kozlov S, Farooqi AA, Naqi A, Lavin M, Khanna KK (2011) ATM protein kinase: the linchpin of cellular defenses to stress. Cell Mol Life Sci 68: 2977–3006

6. Biddlestone-Thorpe L, Sajjad M, Rosenberg E, Beckta JM, Valerie NC, Tokarz M, Adams BR, Wagner AF, Khalil A, Gilfor D et al (2013) ATM kinase inhibition preferentially sensitizes p53-mutant glioma to ionizing radiation. Clin Cancer Res 19: 3189–3200

7. Borok Z, Harboe-Schmidt JE, Brody SL, You Y, Zhou B, Li X, Cannon PM, Kim KJ, Crandall ED, Kasahara N (2001) Vesicular stomatitis virus G-pseudotyped lentivirus vectors mediate efficient apical transduction of polarized quiescent primary alveolar epithelial cells. J Virol 75: 11747–11754

8. Cai X, Chiu YH, Chen ZJ (2014) The cGAS-cGAMP-STING pathway of cytosolic DNA sensing and signaling. Mol Cell 54: 289–296

9. Cao L, Kim S, Xiao C, Wang RH, Coumoul X, Wang X, Li WM, Xu XL, De Soto JA, Takai H et al (2006) ATM-Chk2-p53 activation prevents tumorigenesis at an expense of organ homeostasis upon Brca1 deficiency. Embo j 25: 2167–2177

10. Chan YK, Gack MU (2016) Viral evasion of intracellular DNA and RNA sensing. Nature Reviews Microbiology 14: 360–373

11. Chang LJ, Urlacher V, Iwakuma T, Cui Y, Zucali J (1999) Efficacy and safety analyses of a recombinant human immunodeficiency virus type 1 derived vector system. Gene Ther 6: 715–728

12. Chen G, Shaw MH, Kim YG, Nuñez G (2009) NOD-like receptors: role in innate immunity and inflammatory disease. Annu Rev Pathol 4: 365–398

13. Chen P, Bornhorst J, Aschner M (2018) Manganese metabolism in humans. Front Biosci (Landmark Ed) 23: 1655–1679

14. Connor RI, Chen BK, Choe S, Landau NR (1995) Vpr is required for efficient replication of human immunodeficiency virus type-1 in mononuclear phagocytes. Virology 206: 935–944

15. Cronin J, Zhang XY, Reiser J (2005) Altering the tropism of lentiviral vectors through pseudotyping. Curr Gene Ther 5: 387–398

16. Dai L, Lidie KB, Chen Q, Adelsberger JW, Zheng X, Huang D, Yang J, Lempicki RA, Rehman T, Dewar RL et al (2013) IL-27 inhibits HIV-1 infection in human macrophages by down-regulating host factor SPTBN1 during monocyte to macrophage differentiation. J Exp Med 210: 517–534

17. Dalskov L, Gad HH, Hartmann R (2023) Viral recognition and the antiviral interferon response. Embo j 42: e112907

18. Deguine J, Barton GM (2014) MyD88: a central player in innate immune signaling. F1000Prime Rep 6: 97

19. Delneste Y, Beauvillain C, Jeannin P (2007) [Innate immunity: structure and function of TLRs]. Med Sci (Paris) 23: 67–73

20. Du F, Zhang M, Li X, Yang C, Meng H, Wang D, Chang S, Xu Y, Price B, Sun Y (2014) Dimer monomer transition and dimer re-formation play important role for ATM cellular function during DNA repair. Biochem Biophys Res Commun 452: 1034–1039

21. Fang Z, Jiang W, Liu P, Xia N, Li S, Gu Y (2025) Targeting HIV-1 immune escape mechanisms: Key advances and challenges in HIV-1 vaccine design. Microbiological Research 299: 128229

22. Ferguson BJ, Mansur DS, Peters NE, Ren H, Smith GL (2012) DNA-PK is a DNA sensor for IRF-3-dependent innate immunity. eLife 1: e00047

23. Filipov NM, Seegal RF, Lawrence DA (2005) Manganese potentiates in vitro production of proinflammatory cytokines and nitric oxide by microglia through a nuclear factor kappa B-dependent mechanism. Toxicological Sciences 84: 139–148

24. Franchi L, Warner N, Viani K, Nuñez G (2009) Function of Nod-like receptors in microbial recognition and host defense. Immunol Rev 227: 106–128

25. Gately DP, Hittle JC, Chan GKT, Yen TJ (1998) Characterization of ATM Expression, Localization, and Associated DNA-dependent Protein Kinase Activity. Molecular Biology of the Cell 9: 2361–2374

26. Geijtenbeek TBH, Gringhuis SI (2009) Signalling through C-type lectin receptors: shaping immune responses. Nature Reviews Immunology 9: 465–479

27. Golding SE, Rosenberg E, Valerie N, Hussaini I, Frigerio M, Cockcroft XF, Chong WY, Hummersone M, Rigoreau L, Menear KA et al (2009) Improved ATM kinase inhibitor KU-60019 radiosensitizes glioma cells, compromises insulin, AKT and ERK prosurvival signaling, and inhibits migration and invasion. Mol Cancer Ther 8: 2894–2902

28. He J, Choe S, Walker R, Di Marzio P, Morgan DO, Landau NR (1995) Human immunodeficiency virus type 1 viral protein R (Vpr) arrests cells in the G2 phase of the cell cycle by inhibiting p34cdc2 activity. J Virol 69: 6705–6711

29. Hristova DB, Oliveira M, Wagner E, Melcher A, Harrington KJ, Belot A, Ferguson BJ (2024) DNA-PKcs is required for cGAS/STING-dependent viral DNA sensing in human cells. iScience 27

30. Hu L, Zhang Q (2024) Mechanism of TBK1 activation in cancer cells. Cell Insight 3: 100197

31. Huang H, Kong W, Jean M, Fiches G, Zhou D, Hayashi T, Que J, Santoso N, Zhu J (2019) A CRISPR/Cas9 screen identifies the histone demethylase MINA53 as a novel HIV-1 latency-promoting gene (LPG). Nucleic Acids Research 47: 7333–7347

32. Imamichi T, Chen Q, Sowrirajan B, Yang J, Laverdure S, Marquez M, Mele AR, Watkins C, Adelsberger JW, Higgins J, Sui H (2023) Interleukin-27-induced HIV-resistant dendritic cells suppress reveres transcription following virus entry in an SPTBN1, autophagy, and YB-1 independent manner. PLOS ONE 18: e0287829

33. Ishikawa H, Barber GN (2011) The STING pathway and regulation of innate immune signaling in response to DNA pathogens. Cell Mol Life Sci 68: 1157–1165

34. Iwanaszko M, Kimmel M (2015) NF-κB and IRF pathways: cross-regulation on target genes promoter level. BMC Genomics 16: 307

35. Kane M, Zang TM, Rihn SJ, Zhang F, Kueck T, Alim M, Schoggins J, Rice CM, Wilson SJ, Bieniasz PD (2016) Identification of Interferon-Stimulated Genes with Antiretroviral Activity. Cell Host & Microbe 20: 392–405

36. Karakostis K, Malbert-Colas L, Thermou A, Vojtesek B, Fåhraeus R (2024) The DNA damage sensor ATM kinase interacts with the p53 mRNA and guides the DNA damage response pathway. Molecular Cancer 23: 21

37. Katze MG, He Y, Gale M (2002) Viruses and interferon: a fight for supremacy. Nature Reviews Immunology 2: 675–687

38. König R, Zhou Y, Elleder D, Diamond TL, Bonamy GMC, Irelan JT, Chiang C-y, Tu BP, De Jesus PD, Lilley CE et al (2008) Global Analysis of Host-Pathogen Interactions that Regulate Early-Stage HIV-1 Replication. Cell 135: 49–60

39. Krasnopolsky S, Kuzmina A, Taube R (2020) Genome-wide CRISPR knockout screen identifies ZNF304 as a silencer of HIV transcription that promotes viral latency. PLOS Pathogens 16: e1008834

40. Kumar H, Kawai T, Kato H, Sato S, Takahashi K, Coban C, Yamamoto M, Uematsu S, Ishii KJ, Takeuchi O, Akira S (2006) Essential role of IPS-1 in innate immune responses against RNA viruses. Journal of Experimental Medicine 203: 1795–1803

41. Larabi A, Devos Juliette M, Ng S-L, Nanao Max H, Round A, Maniatis T, Panne D (2013) Crystal Structure and Mechanism of Activation of TANK-Binding Kinase 1. Cell Reports 3: 734–746

42. Lau WC, Li Y, Liu Z, Gao Y, Zhang Q, Huen MS (2016) Structure of the human dimeric ATM kinase. Cell Cycle 15: 1117–1124

43. Li L, Yang X (2018) The Essential Element Manganese, Oxidative Stress, and Metabolic Diseases: Links and Interactions. Oxid Med Cell Longev 2018: 7580707

44. Li Z, Hajian C, Greene WC (2020) Identification of unrecognized host factors promoting HIV-1 latency. PLOS Pathogens 16: e1009055

45. Livak KJ, Schmittgen TD (2001) Analysis of Relative Gene Expression Data Using Real-Time Quantitative PCR and the 2−ΔΔCT Method. Methods 25: 402–408

46. Louis C, Burns C, Wicks I (2018) TANK-Binding Kinase 1-Dependent Responses in Health and Autoimmunity. Frontiers in Immunology Volume 9 – 2018

47. Lv M, Chen M, Zhang R, Zhang W, Wang C, Zhang Y, Wei X, Guan Y, Liu J, Feng K et al (2020) Manganese is critical for antitumor immune responses via cGAS-STING and improves the efficacy of clinical immunotherapy. Cell Research 30: 966–979

48. Ma X, He C, Wang Y, Cao X, Jin Z, Ge Y, Cao Z, An M, Hao L (2025) Mechanisms and Applications of Manganese-Based Nanomaterials in Tumor Diagnosis and Therapy. Biomaterials Research 29: 0158

49. Medzhitov R, Janeway C, Jr. (2000) Innate immunity. N Engl J Med 343: 338–344

50. Medzhitov R, Preston-Hurlburt P, Kopp E, Stadlen A, Chen C, Ghosh S, Janeway CA, Jr. (1998) MyD88 is an adaptor protein in the hToll/IL-1 receptor family signaling pathways. Mol Cell 2: 253–258

51. Ming Q, Liu J, Lv Z, Wang T, Fan R, Zhang Y, Chen M, Sun Y, Han W, Mei Q (2024) Manganese boosts natural killer cell function via cGAS-STING mediated UTX expression. MedComm (2020) 5: e683

52. Mouzakis A, Petrakis V, Tryfonopoulou E, Panopoulou M, Panagopoulos P, Chlichlia K (2025) Mechanisms of Immune Evasion in HIV-1: The Role of Virus-Host Protein Interactions. Curr Issues Mol Biol 47

53. OhAinle M, Helms L, Vermeire J, Roesch F, Humes D, Basom R, Delrow JJ, Overbaugh J, Emerman M (2018) A virus-packageable CRISPR screen identifies host factors mediating interferon inhibition of HIV. eLife 7: e39823

54. Park RJ, Wang T, Koundakjian D, Hultquist JF, Lamothe-Molina P, Monel B, Schumann K, Yu H, Krupzcak KM, Garcia-Beltran W et al (2017) A genome-wide CRISPR screen identifies a restricted set of HIV host dependency factors. Nature Genetics 49: 193–203

55. Paul S, Biswas SR, Milner JP, Tomsick PL, Pickrell AM (2025) Adaptor-Mediated Trafficking of Tank Binding Kinase 1 During Diverse Cellular Processes. Traffic 26: e70000

56. Pham AM, TenOever BR (2010) The IKK Kinases: Operators of Antiviral Signaling. Viruses 2: 55–72

57. Phan LM, Rezaeian AH (2021) ATM: Main Features, Signaling Pathways, and Its Diverse Roles in DNA Damage Response, Tumor Suppression, and Cancer Development. Genes (Basel) 12

58. Pradeu T, Thomma BPHJ, Girardin SE, Lemaitre B (2024) The conceptual foundations of innate immunity: Taking stock 30 years later. Immunity 57: 613–631

59. Rehwinkel J, Gack MU (2020) RIG-I-like receptors: their regulation and roles in RNA sensing. Nature Reviews Immunology 20: 537–551

60. Reis e Sousa C, Yamasaki S, Brown GD (2024) Myeloid C-type lectin receptors in innate immune recognition. Immunity 57: 700–717

61. Roudaire T, Héloir MC, Wendehenne D, Zadoroznyj A, Dubrez L, Poinssot B (2020) Cross Kingdom Immunity: The Role of Immune Receptors and Downstream Signaling in Animal and Plant Cell Death. Front Immunol 11: 612452

62. Runde AP, Mack R, S JP, Zhang J (2022) The role of TBK1 in cancer pathogenesis and anticancer immunity. J Exp Clin Cancer Res 41: 135

63. Sato M, Suemori H, Hata N, Asagiri M, Ogasawara K, Nakao K, Nakaya T, Katsuki M, Noguchi S, Tanaka N, Taniguchi T (2000) Distinct and Essential Roles of Transcription Factors IRF-3 and IRF-7 in Response to Viruses for IFN-α/β Gene Induction. Immunity 13: 539–548

64. Seth RB, Sun L, Ea C-K, Chen ZJ (2005) Identification and Characterization of MAVS, a Mitochondrial Antiviral Signaling Protein that Activates NF-κB and IRF3. Cell 122: 669–682

65. Sharma M, Wagh P, Shinde T, Trimbake D, Tripathy AS (2025) Exploring the Role of Pattern Recognition Receptors as Immunostimulatory Molecules. Immun Inflamm Dis 13: e70150

66. Shibata A, Jeggo PA (2021) ATM’s Role in the Repair of DNA Double-Strand Breaks. Genes (Basel) 12

67. Shu J, Wang X, Yang X, Zhao G (2023) ATM inhibitor KU60019 synergistically sensitizes lung cancer cells to topoisomerase II poisons by multiple mechanisms. Scientific Reports 13: 882

68. So S, Davis AJ, Chen DJ (2009) Autophosphorylation at serine 1981 stabilizes ATM at DNA damage sites. J Cell Biol 187: 977–990

69. Stracker TH, Roig I, Knobel PA, Marjanović M (2013) The ATM signaling network in development and disease. Front Genet 4: 37

70. Sui H, Chen Q, Yang J, Srirattanapirom S, Imamichi T (2022) Manganese enhances DNA-or RNA-mediated innate immune response by inducing phosphorylation of TANK-binding kinase 1. iScience 25: 105352

71. Sui H, Hao M, Chang W, Imamichi T (2021) The Role of Ku70 as a Cytosolic DNA Sensor in Innate Immunity and Beyond. Front Cell Infect Microbiol 11: 761983

72. Sui H, Zhou M, Chen Q, Lane HC, Imamichi T (2013) siRNA enhances DNA-mediated interferon lambda-1 response through crosstalk between RIG-I and IFI16 signalling pathway. Nucleic Acids Research 42: 583–598

73. Sui H, Zhou M, Imamichi H, Jiao X, Sherman BT, Lane HC, Imamichi T (2017) STING is an essential mediator of the Ku70-mediated production of IFN-λ1 in response to exogenous DNA. Science Signaling 10: eaah5054

74. Sun L, Wu J, Du F, Chen X, Chen ZJ (2013) Cyclic GMP-AMP synthase is a cytosolic DNA sensor that activates the type I interferon pathway. Science 339: 786–791

75. Sun S, Xu Y, Qiu M, Jiang S, Cao Q, Luo J, Zhang T, Chen N, Zheng W, Meurens F et al (2023) Manganese Mediates Its Antiviral Functions in a cGAS-STING Pathway Independent Manner. Viruses 15

76. Suzuki T, Oshiumi H, Miyashita M, Aly HH, Matsumoto M, Seya T (2013) Cell type-specific subcellular localization of phospho-TBK1 in response to cytoplasmic viral DNA. PLoS One 8: e83639

77. Taylor JP, Cash MN, Santostefano KE, Nakanishi M, Terada N, Wallet MA (2018) CRISPR/Cas9 knockout of USP18 enhances type I IFN responsiveness and restricts HIV-1 infection in macrophages. Journal of Leukocyte Biology 103: 1225–1240

78. tenOever Benjamin R, Sharma S, Zou W, Sun Q, Grandvaux N, Julkunen I, Hemmi H, Yamamoto M, Akira S, Yeh W-C et al (2004) Activation of TBK1 and IKKε Kinases by Vesicular Stomatitis Virus Infection and the Role of Viral Ribonucleoprotein in the Development of Interferon Antiviral Immunity. Journal of Virology 78: 10636–10649

79. Thompson MR, Kaminski JJ, Kurt-Jones EA, Fitzgerald KA (2011) Pattern recognition receptors and the innate immune response to viral infection. Viruses 3: 920–940

80. Tidball AM, Bryan MR, Uhouse MA, Kumar KK, Aboud AA, Feist JE, Ess KC, Neely MD, Aschner M, Bowman AB (2015) A novel manganese-dependent ATM-p53 signaling pathway is selectively impaired in patient-based neuroprogenitor and murine striatal models of Huntington’s disease. Hum Mol Genet 24: 1929–1944

81. Ueno S, Sudo T, Hirasawa A (2022) ATM: Functions of ATM Kinase and Its Relevance to Hereditary Tumors. International Journal of Molecular Sciences 23: 523

82. Unterholzner L, Keating SE, Baran M, Horan KA, Jensen SB, Sharma S, Sirois CM, Jin T, Latz E, Xiao TS et al (2010) IFI16 is an innate immune sensor for intracellular DNA. Nat Immunol 11: 997–1004

83. Wang B, Zhang F, Wu X, Ji M (2024) TBK1 is paradoxical in tumor development: a focus on the pathway mediating IFN-I expression. Frontiers in Immunology Volume 15–2024

84. Wang C, Guan Y, Lv M, Zhang R, Guo Z, Wei X, Du X, Yang J, Li T, Wan Y et al (2018) Manganese Increases the Sensitivity of the cGAS-STING Pathway for Double-Stranded DNA and Is Required for the Host Defense against DNA Viruses. Immunity 48: 675–687.e677

85. Warren C, Pavletich NP (2022) Structure of the human ATM kinase and mechanism of Nbs1 binding. Elife 11

86. Weiss WA, Taylor SS, Shokat KM (2007) Recognizing and exploiting differences between RNAi and small-molecule inhibitors. Nat Chem Biol 3: 739–744

87. West AP, Koblansky AA, Ghosh S (2006) Recognition and signaling by toll-like receptors. Annu Rev Cell Dev Biol 22: 409–437

88. Xu L-G, Wang Y-Y, Han K-J, Li L-Y, Zhai Z, Shu H-B (2005) VISA Is an Adapter Protein Required for Virus-Triggered IFN-β Signaling. Molecular Cell 19: 727–740

89. Yu L, Liu P (2021) Cytosolic DNA sensing by cGAS: regulation, function, and human diseases. Signal Transduction and Targeted Therapy 6: 170

90. Zhang Q, Green MD, Lang X, Lazarus J, Parsels JD, Wei S, Parsels LA, Shi J, Ramnath N, Wahl DR et al (2019) Inhibition of ATM Increases Interferon Signaling and Sensitizes Pancreatic Cancer to Immune Checkpoint Blockade Therapy. Cancer Res 79: 3940–3951

91. Zhang R, Wang C, Guan Y, Wei X, Sha M, Yi M, Jing M, Lv M, Guo W, Xu J et al (2021) Manganese salts function as potent adjuvants. Cell Mol Immunol 18: 1222–1234

92. Zhang X, Brann TW, Zhou M, Yang J, Oguariri RM, Lidie KB, Imamichi H, Huang DW, Lempicki RA, Baseler MW et al (2011) Cutting edge: Ku70 is a novel cytosolic DNA sensor that induces type III rather than type I IFN. J Immunol 186: 4541–4545

93. Zhou H, Xu M, Huang Q, Gates AT, Zhang XD, Castle JC, Stec E, Ferrer M, Strulovici B, Hazuda DJ, Espeseth AS (2008) Genome-Scale RNAi Screen for Host Factors Required for HIV Replication. Cell Host & Microbe 4: 495–504

94. Zhou Y, Zhou B, Pache L, Chang M, Khodabakhshi AH, Tanaseichuk O, Benner C, Chanda SK (2019) Metascape provides a biologist-oriented resource for the analysis of systems-level datasets. Nat Commun 10: 1523

